# Cycling with blood flow restriction improves performance and muscle K^+^ handling and blunts the effect of antioxidant infusion in humans

**DOI:** 10.1101/375881

**Authors:** Danny Christiansen, Kasper H. Eibye, Villads Rasmussen, Hans M. Voldbye, Martin Thomassen, Michael Nyberg, Thomas G.P. Gunnarsson, Casper Skovgaard, Mads S. Lindskrog, David J. Bishop, Morten Hostrup, Jens Bangsbo

## Abstract

We examined if blood flow restriction (BFR) would augment training-induced improvements in muscle K^+^ handling and performance during intense exercise in men, and if these adaptations would be associated with an effect of muscle antioxidant function on thigh K^+^ release and with fibre type-dependent modulation of Na^+^,K^+^-ATPase-isoform abundance and FXYD1 phosphorylation. Ten recreationally-active men (25 ± 4 y, 49.7 ± 5.3 mL∙kg^-1^∙min^-1^) performed 6 weeks of interval cycling, where one leg trained without (control; CON-leg) and the other leg with BFR (BFR-leg, pressure: 178 mmHg). Before and after training, catheters were inserted into the femoral artery and vein, and blood flow was assessed during single-leg knee-extensions at 25% (Ex1) and 90% of leg peak aerobic power (Ex2) with intravenous infusion of N-acetylcysteine (NAC) or saline (placebo), and a resting muscle biopsy was collected. After training, performance during exhaustive exercise increased to a greater extent in BFR-leg (23%) than in CON-leg (12%, p<0.05), whereas thigh K^+^ release during Ex2 was attenuated in BFR-leg only (p<0.05). Before training, NAC depressed K^+^ release during Ex1 (p<0.05), but not during Ex2 (p>0.05). After training, this effect was blunted in BFR-leg (p<0.05), whilst the abundance of Na^+^,K^+^-ATPase-isoform α_1_ in type-II (51%), β_1_ in type-I (33%), and FXYD1 in type-I (108%) and type-II (60%) fibres was higher in BFR-leg (p<0.05; vs. CON-leg). Thus, interval training with BFR elicits greater improvements in performance and reduces muscle net K^+^ release during intense exercise, which may be caused by elevated ROS scavenging and fibre type-dependent increases in Na^+^,K^+^-ATPase-isoform abundance.

**Key points:**

- Here, we provide evidence that reactive oxygen species (ROS) play a role in regulating K^+^ homeostasis in the untrained musculature of humans, as indicated by attenuated thigh K^+^ efflux during exercise with concomitant antioxidant infusion.
- We also demonstrate that interval training with blood flow restriction (BFR) augments improvements in performance and reduces K^+^ release from contracting muscles during intense exercise
- The effect of training with BFR on muscle K^+^ handling appears to be partly mediated by increasing the protection against ROS, since the effect of antioxidant infusion was blunted after training with restricted blood flow.
- Further, training with BFR resulted in higher abundance of Na^+^,K^+^-ATPase-isoform α_1_ in type-II (51%), β_1_ in type-I (33%), and FXYD1 in type-I (108%) and type-II (60%) muscle fibres. This suggests fibre type-specific adaptations in Na^+^,K^+^-ATPase-isoform content are also important for improvements in muscle K^+^ handling by training with BFR in humans.

## Introduction

Training with blood flow restriction (BFR) has become a well-recognised strategy to facilitate muscle hypertrophy and strength (Pearson & Hussain, 2015), and there is some indications that BFR may augment training adaptations in muscle oxidative capacity (Sundberg *et al.*, 1993; Christiansen *et al.*, 2017a; Paton *et al.*, 2017) and exercise performance (Sundberg *et al.*, 1993; Manimmanakorn *et al.*, 2013). BFR is often achieved by inflation of an occlusion cuff around the exercising limb(s), resulting in a severely hypoxic muscular environment and an increased rate of anaerobic glycolytic flux (Corvino *et al.*, 2017; Christiansen *et al.*, 2018). Subsequent deflation of the cuff raises muscle oxygen delivery (Gundermann *et al.*, 2012). Together, these mechanisms create favourable conditions for ROS production (Clanton, 2007; Stoner *et al.*, 2007) and ionic (K^+^ and Ca^2+^) perturbations (Shimoda & Polak, 2011), both of which are potent stimuli for increasing the expression of K^+^ regulatory systems, such as the Na^+^, K^+^-ATPase (Murphy *et al.*, 2006; Silva & Soares-da-Silva, 2007). Thus, training with BFR may be a potent strategy to promote adaptations in skeletal muscle K^+^ handling.

Studies in intact muscle fibres have demonstrated that reactive oxygen species (ROS) generated during muscle contractions, or exogenously administered, modulate muscle force production in both a time and dose-dependent manner (Reid *et al.*, 1993). At high concentrations, ROS retard muscle force development, which has been partially ascribed to disruption of myocellular K^+^ homeostasis (Cerbai *et al.*, 1991). This is consistent with observations that Na^+^,K^+^-ATPase activity is depressed by ROS exposure in animals (Glynn, 1963; Boldyrev & Kurella, 1996), or elevated during exercise by antioxidant treatment in humans (McKenna *et al.*, 2006).

Antioxidant treatment has been shown to attenuate the rise in arterialised-venous plasma K^+^ concentrations during submaximal exercise (McKenna *et al.*, 2006), suggesting ROS may perturb whole-body K^+^ homeostasis. However, systemic K^+^ levels may inaccurately reflect locomotor muscle K^+^ homeostasis (Nielsen *et al.*, 2004; Gunnarsson *et al.*, 2013), indicating the need to directly assess K^+^ efflux from exercising musculature to elucidate the role of ROS in regulating K^+^ homeostasis in human skeletal muscle. In humans, intense training enhances K^+^ handling (Nielsen *et al.*, 2004) and ROS defence in skeletal muscle (Hellsten *et al.*, 1996; Leeuwenburgh *et al.*, 1997). Based on these findings and the *in vitro* studies described above, there is rationale to believe that training-induced improvements in muscle K^+^ handling are associated with enhanced protection against oxidative damage. However, this has not been examined in humans.

In rats, excitation-induced fatigue manifests earlier in fast-twitch than in slow-twitch muscle, which has been linked with faster accumulation of extracellular K^+^ and resultant inexcitability (Clausen *et al.*, 2004). These differences have partly been ascribed to a different expression of Na^+^,K^+^-ATPase isoforms (α_1_, α_2_, β_1_) between fibre types (Fowles *et al.*, 2004). Thus, recent observations in humans show that Na^+^,K^+^-ATPase isoforms are expressed in a fibre type-specific manner and that their content is differently altered by training (Thomassen *et al.*, 2013; Wyckelsma *et al.*, 2015; Christiansen *et al.*, 2017b). Despite these observations, it remains unresolved how training-induced improvements in muscle K^+^ handling are associated with modulation of Na^+^,K^+^-ATPase-isoform content at the fibre-type level in humans. Further, in myocytes, raising FXYD content counteracts Na^+^,K^+^-ATPase dysfunction induced by oxidative damage (Bibert *et al.*, 2011), whereas elevated FXYD1 phosphorylation increases Na^+^ affinity and ouabain-sensitive activity of the Na^+^,K^+^-ATPase (Reis *et al.*, 2005; Ingwersen *et al.*, 2011). Therefore, improvements in muscle K^+^ handling may also be obtained by increasing FXYD1 abundance, phosphorylation, or both. However, to our knowledge, no studies in humans have explored the effects of training on the relationship between FXYD1 content/phosphorylation and muscle K^+^ handling during exercise. Because FXYD1 has been shown to be either expressed or phosphorylated in a fibre type-specific manner in sedentary rats (Reis *et al.*, 2005; Juel, 2009; Thomassen *et al.*, 2011) and recreationally active humans (Thomassen *et al.*, 2013; Christiansen *et al.*, 2017b), this relationship should be studied at the fibre-type level.

Thus, the primary aims of the present study were to examine the effects of training with BFR on performance and leg K^+^ handling during exercise in men, and to elucidate the muscular mechanisms involved. The following hypotheses were tested: 1) Training with BFR elicits greater improvements in exercise performance and attenuates thigh K^+^ release during exercise compared to training without BFR. 2) Antioxidant infusion counteracts an increase in thigh K^+^ release during exercise 3) Training with BFR blunts the effect of antioxidant infusion on thigh K^+^ release during exercise. 4) Improvements in leg K^+^ handling induced by training with BFR are associated with fibre type-dependent modulation of Na^+^,K^+^-ATPase isoform (α_1_, α_2_, β_1_, FXYD1) abundance and FXYD1 phosphorylation.

## Methods

### Ethical Approval

Participants were informed about the requirements, benefits, and potential risks of the study before providing informed written consent to participate. The experimental procedures adhered to the standards in the latest revision of the Declaration of Helsinki and were approved by the Human Research Ethics Committee of the Capital Region of Denmark (approval no. H-16000377).

### Participants

Thirteen recreationally active men (mean ± SD: age: 25 ± 4 y; height: 183 ± 6 cm; body mass: 83.6 ± 14 kg; VO_2max_: 49.7 ± 5.3 mL∙kg^-1^∙min^-1^) volunteered to participate in the study. They were non-smokers, free of medication, and engaged in regular physical activity (i.e. soccer, fitness/gym, swimming or running) 1-3 times per week. The participants did not use anti-inflammatory drugs or supplements and were asked to maintain their routine physical activity throughout the study. Two participants withdrew after the first experimental day due to the invasive nature of the experiment, whereas one participant dropped out two weeks into the familiarisation period due to loss of motivation. Thus, ten men completed the study.

### Experimental design

This study used an intra-subject design. The participants’ legs were randomly assigned in a counterbalanced fashion to a 6-wk training intervention consisting of interval cycling with BFR for one leg (BFR-leg) and without BFR for the other leg (CON-leg). Randomisation took into account the dominant (i.e. preferred kicking) leg and was performed using a random-number generator in Excel (Ver. 2013, MS Office, Microsoft, Washington, USA). The legs were trained simultaneously and their force output matched during every training session.

Before and after the training period, the performance of each leg was assessed during incremental knee-extensor exercise to exhaustion in a Krogh ergometer, allowing isolated work with the quadriceps muscles (Andersen & Saltin, 1985). After at least 48 h, participants completed an experimental day (detailed later), on which femoral arterial and venous blood K^+^ concentrations were measured, and femoral arterial blood flow assessed, in each leg to determine thigh K^+^ release during exercise at various intensity under intravenous infusion of antioxidant (N-acetylcysteine) or placebo (saline). A resting muscle biopsy was also obtained from each leg to determine Na^+^,K^+^-ATPase-isoform abundance and FXYD1 phosphorylation in type I and II fibres. In addition, participants performed an incremental exercise test to exhaustion on a bike ergometer to determine training intensity and maximal oxygen consumption (VO_2max_). To minimise any placebo effect, participants were not informed about which intervention was hypothesised to be the most beneficial for their performance and muscle response. All tests were performed under standard laboratory conditions (~23 ºC, ~33 % humidity) in the Department of Nutrition, Exercise and Sport (University of Copenhagen, Denmark). Participants were asked to abstain from alcohol, caffeine, and strenuous physical activity for 24 h prior to each laboratory visit and recorded their dietary pattern in this period. They were asked to replicate this pattern before each visit.

### Incremental knee-extensor exercise test

On the day of the first incremental, single-leg knee-extensor exercise test, participants were instructed to consume a light meal 3 h prior to the start of the test and to replicate this dietary pattern prior to each subsequent non-invasive test (i.e. without sampling of blood and muscle tissue). In the 3 h before and during these tests, only water was allowed *ad libitum*. Upon arrival at the laboratory, the participants changed clothes, were asked about their sleep, food and liquid intake, and their body mass was recorded. Next, a 5-min warm-up was performed at 100 W (75 rpm) on a cycle ergometer (Monark Exercise, AB, Vansbro, Sweden). The participants were then placed in the knee-extensor model. After 15 min of rest, the participants carried out the test, whilst sitting upright with a hip angle of ~120°. The test commenced with 5 min at 24 W, followed by an increase in workload of 6 W per min until volitional exhaustion, defined as an inability to sustain the required workload at a kicking frequency of 60 rpm. Each leg was tested separately in a randomised order interspersed by at least 30 min of rest. This order was kept the same for each participant before and after the training period. The power output attained at task failure (iPPO) was calculated as the sum of the power output at the last completed stage plus the increment (6 W) multiplied by the fractional time spend at the last (incomplete) stage. The first execution of the test before the familiarisation was used to establish an exercise intensity that would elicit task failure after 3-9 min. This relative intensity was used in Ex2 on the experimental day. To enable calculation of the workload to be used in Ex1, this test was always executed prior to the experimental day before and after the training period.

### Incremental bike ergometer test

Participants performed the incremental cycling test on a bike ergometer (Monark Exercise, AB, Vansbro, Sweden) using cycling shoes with cleats. Before the test, participants were instrumented with a facemask covering the mouth and nose, which was connected to an online gas-analysing system (Oxycon Pro, Jaeger, Germany) to measure (breath by breath) inspired and expired gasses. The test comprised of repeated 4-min cycling bouts interspersed by 1 min of rest. The first bout commenced at 90 W, after which the workload was increased by 30 W at the onset of each subsequent bout until volitional exhaustion, defined as an inability to maintain the required workload. After 5 min of additional rest, the participants commenced exercising for 1 min at the workload of the last completed bout, after which the workload was increased by 10 W per min until exhaustion to ascertain a plateau in VO_2max_. VO_2max_ was determined as the maximum 15-s plateau in oxygen uptake during the test. The maximum workload (W_max_) was calculated as the sum of the workload of the last completed bout plus the increment (30 W) multiplied by the fractional time spend on the last (incomplete) bout.

### Training

Training sessions took place indoor on a Tomahawk IC7 cycle ergometer (Indoor Cycling Group, Nürnberg, Germany) at a local gym (fitnessdk Adelgade, Copenhagen, Denmark). Prior to the training period, participants underwent four weeks of training to become familiarised with the training protocol, and to be accustomed to maintain their preferred cadence (range: 75-85 rpm) and to adjust their workloads. In this period, they completed (mean ± SD) 8 ± 3 training sessions similar to those in the training period, with sessions separated by at least 48 h. During the intervention period, the participants trained three times per week, amounting to a total of 17 ± 1 (mean ± SD) training sessions. Each training session commenced with a 5-min warm-up at 30% of maximum workload (W_max_), determined from the graded exercise test, followed by 2 min of rest. Next, three periods of 3 x 2-min cycling bouts were performed separated by 1 min and each period by 2 min of active recovery (i.e., pedalling freely without workload). The target intensity of the first, second, and third period was 60, 70, and 80% W_max_, respectively. The intensity and cuff pressure was noted, and the participants reported their overall rating of perceived exertion (RPE) on the 6-20 BORG-scale (Borg, 1982). During the intervention period, the training intensity was (mean ± SD) 61 ± 3, 72 ± 4, and 81 ± 10% W_max_ in first, second, and third period, respectively. RPE was 13 ± 1, 16 ± 2, and 19 ± 1, respectively. An investigator provided verbal encouragement to the participants throughout each session.

### Matching of leg force during training

The participants performed each training session with force sensor insoles (Novel GmbH, Munich, Germany) in their cycling shoes to measure the force generated during every pedal rotation. The force output was wirelessly transmitted via Bluetooth to an iPad Mini 2 (Apple Inc. California, USA) using Pedoped software (Pedoped Expert, Novel GmbH, Munich, Germany). The iPad was mounted on the participants’ handlebar to enable visual feedback in real-time about the force generated by each leg. Before use, every insole was validated against the same force plate (Kistler, Germany). The coefficient of variation for the insole signal, relative to the force plate, was ≤ 5 %. Throughout each session, the participants were reminded repeatedly by the investigators to maintain the same peak force between the legs. After each session, the force data was stored on the iPads and later exported to a laptop for further analysis in Excel (Ver. 2013, MS Office, Microsoft, Washington USA). During the intervention period, the force (mean ± SD) generated during each pedal rotation by the CON-leg and BFR-leg was 102 ± 10 and 104 ± 8 N in the first period, 107 ± 9 and 109 ± 11 N in the second period, and 123 ± 15 and 121 ± 13 N in the third period, respectively. There was no difference in force output between the CON-leg and BFR-leg at any intensity (p > 0.05).

### Blood flow restriction during training

In the BFR-leg, the femoral arterial blood flow was reduced during each cycling bout by inflation of a 13-cm wide nylon cuff (Riester, Germany). The cuff was placed around the most proximal part of the leg and was loosely kept in place by an elastic bandage throughout each training session. The cuff was inflated ~10 s prior to and deflated immediately after every exercise bout to induce repeated reductions in muscle blood perfusion during exercise followed by reperfusion recovery, in line with the method used previously (Christiansen *et al.*, 2018). The cuff pressure during training was (mean ± SD) 178 ± 10 mmHg and varied from 155 ± 7 mmHg in the relaxed state to 200 ± 12 mmHg in the contracting state of the quadriceps muscles.

### Assessment of blood flow perturbations induced by BFR

To determine the effects of BFR on femoral arterial blood flow during training, five participants completed an additional experiment during the familiarisation period. In this experiment, they performed two identical knee-extensor exercise bouts at moderate intensity (12W) with one leg to simulate the training stimulus invoked on the knee-extensor muscle group during the intervention period. One bout was performed without (CON) and the other with an occlusion cuff around the leg inflated to ∼ 178 mmHg (as during the intervention period). The bouts were separated by one hour and their order randomised. The leg used for the experiment was randomly chosen based on knowledge about dominant (preferred kicking) leg. During the experiment, femoral arterial blood flow was measured by ultrasound Doppler at rest before and after BFR, at 20s, 40s, 1 min, 2 min, and 2.5 min during exercise, and at 20s, 40s, 1 min, 2 min, and 3 min in recovery.

### Experimental day

Participants arrived at the laboratory at 8 am after consuming a standardised, self-chosen breakfast (e.g., 140 g oats, 1 banana, 50 g raisins, 350 mL skim milk, and 0.75 L water) ~2 h prior to arrival. After 15 min of rest in the supine position, catheters (20 gauge) were inserted into the femoral artery and vein of both legs under local anaesthesia (lidocaine, 20 mg∙mL^−1^) with use of ultrasound Doppler. After 15 min of rest, the participants were seated in the knee-extensor model, where they rested for another 15 to 25 min, before commencing the exercise protocol illustrated in Fig. 1. The protocol comprised of two sets of single-leg exercise with each leg in the knee-extensor model. Each set consisted of exercise at an intensity of 25% iPPO for 10 min (Ex1), followed by 25 min of rest and an intense exercise bout at 90% iPPO to exhaustion (Ex2), which was defined as an inability to sustain the required workload at ≥50 rpm. If the participants were able to sustain exercise longer than 4 min, the workload was increased by 6 W per min after 4 min until exhaustion. Exercise was always started by passive movement of the leg for 10 s (by investigator’s rotation of the flywheel) to reach a cadence of 60 rpm. In the 15 min before and during the first set, saline (placebo; 20 mg∙mL^−1^) was infused at a constant rate (1 mL∙h^−1^) in the femoral vein of the resting leg. Twenty minutes prior to the second set, infusion of the antioxidant N-acetylcysteine (NAC) was commenced at a rate of 125 mg∙kg^−1^∙h^−1^ for 15 min to reach a plateau in plasma NAC concentration, followed by constant infusion at 25 mg∙kg^−1^∙h^−1^ for the remainder of the second set, in line with the NAC infusion protocol previously used (Medved *et al.*, 2003). In this set, Ex2 was terminated at the point of exhaustion reached during the first set with placebo infusion, if the participants did not terminate exercise prior to this time point. Infusions of placebo and NAC were separated by ~45 min of rest. Within the first 10 min of this resting period, participants consumed two pieces of fruit (i.e. an apple, a banana and/or an orange). Except for this meal, only water was allowed *ad libitum* throughout the day. Participants were blinded to the infusions and reported no adverse effects of infusions. The exercise order of the legs was randomised, whereas the order for each participant was the same after relative to before the intervention period. After training, Ex2 was stopped at the time of exhaustion reached in Ex2 before the intervention. Blood was sampled from the femoral artery and vein of the working leg, and femoral arterial blood flow was measured by ultrasound Doppler over the same leg, at rest before and after infusions and at 20 s, 1 min, 3 min, and 8 min of Ex1, and 20 s, 1 min, and 3 min of Ex2, and after 20 s 1 min, 2 min, and 3 min of recovery from each exercise bout. To correct for the mean transit time from artery to vein, venous blood was collected five seconds after each arterial sample (Bangsbo *et al.*, 2000). Leg blood flow was measured continuously during blood sampling. In addition, a muscle biopsy was obtained approximately 10 min prior to Ex2. The mean ± SD workload in Ex1 for CON-leg and BFR-leg was 17 ± 4 and 16 ± 4 W before the intervention period, and 18 ± 3 and 20 ± 3 W after the intervention period, respectively. The mean ± SD duration of Ex2 for CON-leg and BFR-leg was 4:52 ± 1:35 and 4:44 ± 1:07 min with placebo, and 4:37 ± 1:30 and 4:29 ± 1:16 min with NAC infusion at Pre, and 4:50 ± 1:34 and 4:43 ± 1:07 with placebo and 4:37 ± 1:29 and 4:29 ± 1:15 min with NAC infusion at Post, respectively. The workload was the same for placebo vs. NAC infusion.

**Figure 1.**
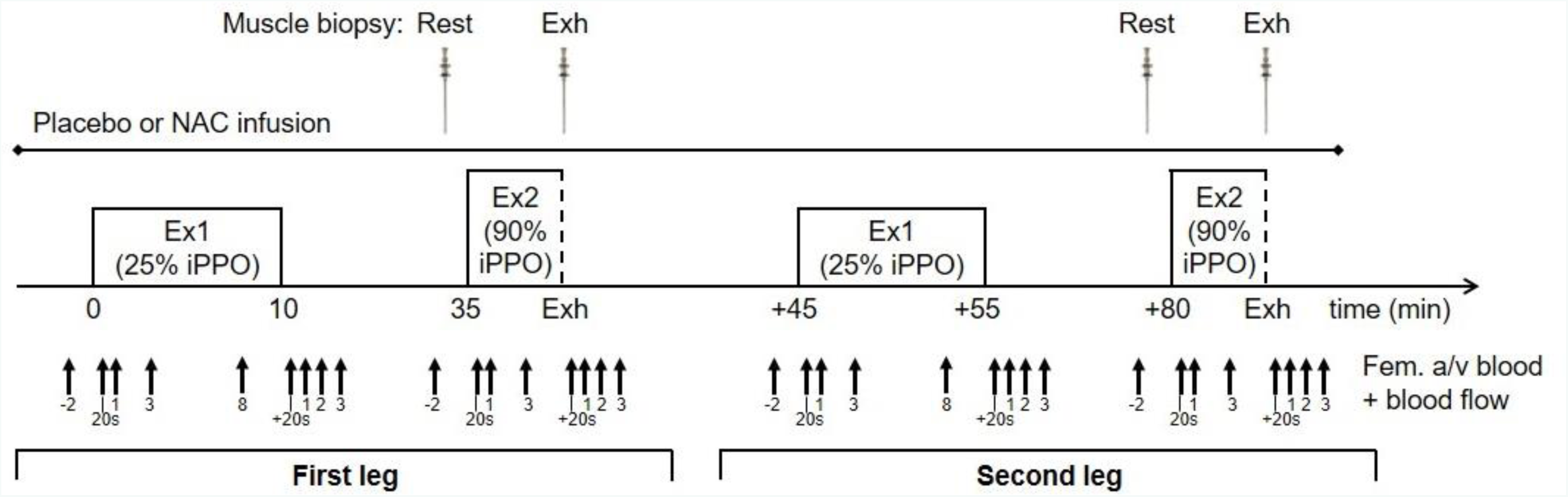
*Illustration of the experimental day*. Participants performed two exercise sets with each leg in the knee-extensor exercise model in a randomised order. Each set consisted of a 10-min exercise bout at 25% incremental peak power output (iPPO; Ex1), followed by an exhaustive bout at 90% of pre-training iPPO (Ex2). During the first set, saline (placebo) was intravenously infused, whereas the antioxidant N-acetylcysteine (NAC) was infused during the second set after 45 min of rest. Blood was sampled from the femoral artery and vein (Fem. a/v blood) of the active leg, and femoral arterial blood flow was measured over the same leg at the time points indicated. A muscle biopsy was obtained at rest before the intervention (Rest).

### Measurement of leg blood flow

Femoral arterial blood flow was measured by ultrasound Doppler (Logic E9, GE Healthcare, Pittsburg, PA, USA) equipped with a linear probe operating at an image frequency of 9 MHz and a Doppler frequency of 3.1 MHz. Blood velocity was measured in the femoral artery distal to the inguinal ligament, but above the bifurcation into the superficial and profound femoral branch. During all measurements, the insonation angle was kept as low as possible and was always below 60 deg. Sample volume was adjusted to include the entire vessel lumen, while avoiding vessel walls. A low-velocity filter (velocities <1.8 m/s) removed noise caused by turbulence at the vascular wall. Doppler tracings and B-mode images were recorded continuously, and Doppler tracings were averaged over 10-16 s at the time of blood sampling. Vessel diameter was determined for each measurement. Artery diameter was measured during systole at rest, and upon peak vessel expansion during exercise and recovery. Diameter was evaluated as the average of three measurements taken perpendicular to the vessel wall at the site of blood velocity measurements.

### Blood sampling

Approximately 2 mL of blood was drawn per sample with a heparinised syringe. Samples were immediately stored on ice until being analysed for K^+^ concentration on an ABL 800 Flex (Radiometer, Copenhagen, Denmark). To ensure collection of circulating blood, 2 mL of blood was withdrawn prior to each sample and reintroduced into the venous circulation after the recovery.

### Muscle biopsy

All biopsies were obtained from the vastus lateralis muscle using the Bergström needle biopsy technique with suction. In preparation, a small incision was made through the skin and muscle fascia under local anaesthesia (2 mL 1% Xylocaine). Incisions were separated by approximately 1-2 cm. Immediately after collection, samples were frozen in liquid nitrogen and later stored at –80 °C until being analysed. The incisions were covered with sterile Band-Aid strips and a Tegaderm film-dressing.

### Dissection and fibre typing of muscle fibre segments

Approximately 15 mg w.w. muscle per sample was freeze-dried for 48 h, after which a minimum of 30 single-fibre segments per sample (range: 30 to 60; total *n* = 2670 fibres) were isolated under a stereo microscope using fine jeweller’s forceps at standard environmental conditions (22ºC, <30 % humidity). After dissection, segments were incubated for 1 h at room temperature in 10-µL denaturing buffer (0.125 M Tris-HCl, 10% glycerol, 4% sodium dodecyl sulphate, 4 M urea, 10% mercaptoethanol, and 0.001% bromophenol blue, pH 6.8) before being stored at −80°C until further analysis.

The fibre type of each fibre segment was determined using a novel dot blotting method described elsewhere (D Christiansen, MJ MacInnis, E Zacharewicz, BP Frankish, H Xu, RM Murphy, *in review in Acta Physiologica*). Two 0.45-µm PVDF membranes were activated for 15 s in 96% ethanol and equilibrated for two minutes in transfer buffer (25 mM Tris, 192 mM glycine, 0.015% SDS, 96% ethanol, pH 8.3). A 1.5-µL aliquot of denatured sample was spotted onto each membrane. The membranes were then placed at room temperature on a dry piece of filter paper to dry completely (10 to 20 min). The membranes were reactivated in ethanol and re-equilibrated in transfer buffer, and quickly rinsed in Tris-buffered saline-Tween (TBST), after which they were blocked for 5 to 45 min in 5% skim milk in TBST (blocking buffer). Next, one membrane was incubated (1 in 200) with myosin heavy chain I (MHCI) antibody, whereas the other membrane was probed (1 in 200) with myosin heavy chain IIa (MHCIIa) antibody, for 2 to 3 h at room temperature with gentle rocking. Antibody details are shown in Table 1. After two quick washes in blocking buffer, the membranes were incubated (1 in 5.000) for 1 h at room temperature with different horse-radish peroxidase (HRP)-conjugated secondary antibodies (MHCI: goat anti-mouse IgM, #62-6820, Thermo Fisher Scientific; MHCIIa: goat anti-mouse IgG, #PIE31430, Thermo Fisher Scientific). After two 2-min washes in TBST, the membranes were incubated with Clarity enhanced chemiluminescence reagent (Bio-Rad, Hercules, CA, USA). Imaging was performed on a ChemiDoc MP (Bio-Rad). The remainder of each denatured segment (7 µL) was organised into one of two groups of fibres (type I and type II) per sample according to MHC expression. The number of segments included in each group of fibres per sample ranged from 5 to 27 for type I and 6 to 23 for type II. Hybrid fibres (expressing both MHC isoforms) and blank fibres (i.e. no protein present or a clean type IIx fibre) were excluded from analysis.

**Table 1.**
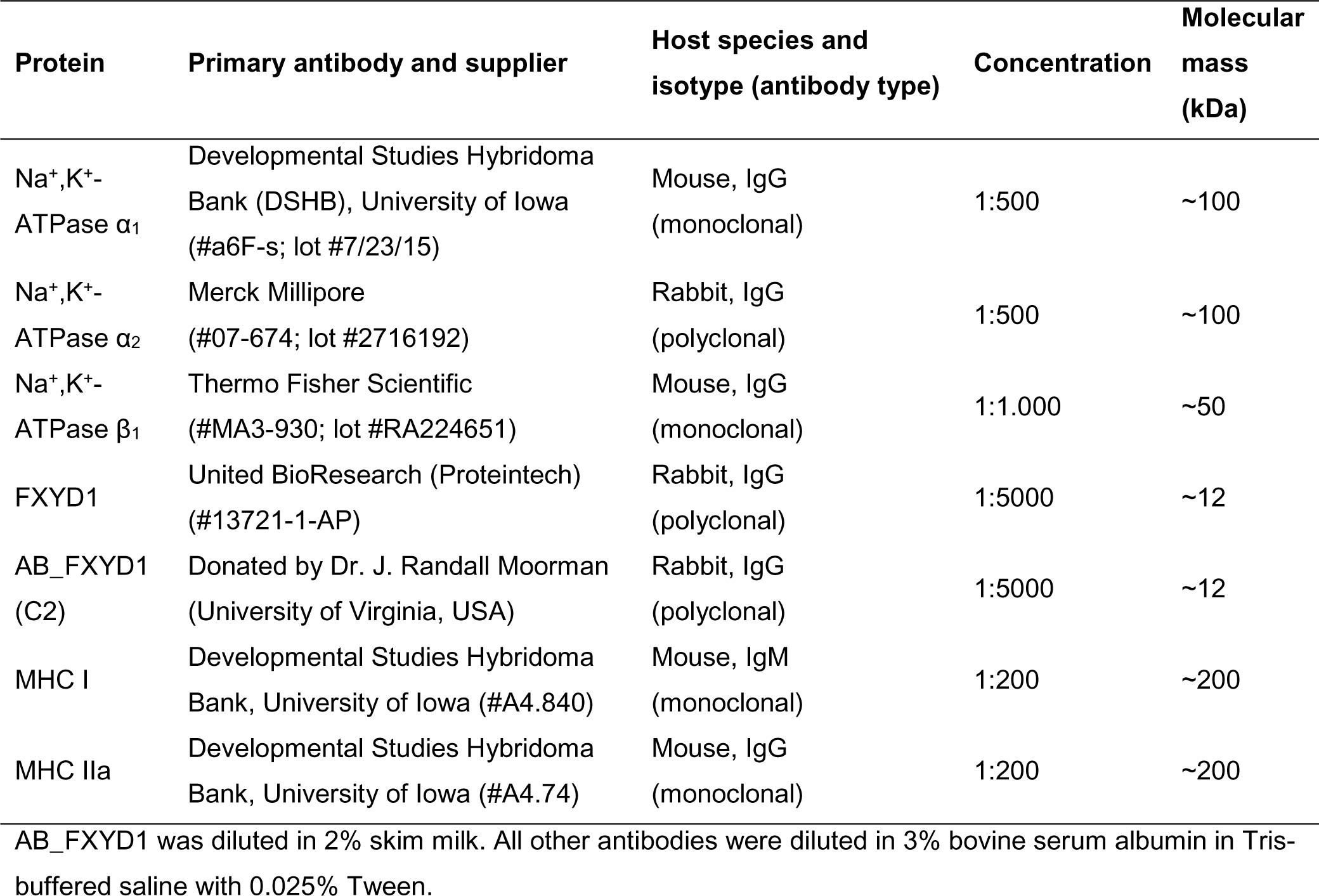
Primary antibodies used for dot blotting and/or western blotting

### Immunoblotting

Protein abundance and phosphorylation were determined by western blotting. Approximately 1 μg protein per type-I and type-II fibre sample were loaded in each well on a 4 to 15% Criterion TGX stain-free gel (Bio-Rad, Hercules, CA). All samples from each participant were loaded onto the same gel, along with two 4-point calibration curves consisting of mixed-fibre human muscle lysate with a known protein concentration (0.25 to 3 μg protein) and two protein ladders (Precision Plus Protein All Blue, #1610373, Bio-Rad). Proteins were separated by electrophoresis (45 min at 200V), after which the gels were UV-activated for 5 min (ChemiDoc, Bio-Rad) to enable stain-free quantification of total protein. Proteins were semi-dried transferred to a 0.45 µm PVDF-membrane for 45 min at 14 V (Amersham TE 70, GE Healthcare) using transfer buffer (25 mM Tris, 190 mM glycine and 20 % methanol). Membranes were blocked for 1 h at room temperature in blocking buffer using gentle rocking. To allow multiple proteins to be quantified on the same membrane, membranes were cut horizontally at the required molecular weights using the two protein ladders as markers before probing with primary antibodies overnight at 4 °C with constant, gentle rocking. Antibody details are presented in Table 1. Primary antibodies were diluted in 3% bovine serum albumin (BSA) in Tris-buffered saline with 0.1% Tween (TBST). After incubation, the membranes were washed in TBST and probed with HRP-conjugated secondary antibody (goat anti-mouse or goat-anti-rabbit immunoglobulins) for 1 h with rocking at room temperature. Protein bands were visualized using enhanced chemiluminescence (SuperSignal West Femto, Rockford, Pierce, IL, USA) on a ChemiDoc MP imaging system (Bio-Rad). Quantification of bands was performed in Image Lab 5.2.1 (Bio-Rad). Protein abundance and phosphorylation in each sample was determined by normalising the density for a given sample to that of the slope of the calibration curve, as well as to the total protein amount in each lane on the stain-free gel image. To improve reliability, the mean of the duplicate readings of a given protein amount from the two calibration curves on the same membrane was used for the analysis. The linearity (r^2^) of calibration curves for total protein and blots was ≥0.98. Further, only first probes were used for analysis. The same blinded researcher was responsible for analysing all western blots included in the study. Based on agreement between two independent researchers, who conducted visual inspections of the blots, fibre pools and points on calibration curves were excluded if their band was unable to be validly quantified due to noise on the image caused by artefacts or if they were too faint or were saturated. This resulted in *n* = 9 for Na^+^,K^+^-ATPase α_1_, *n* = 3 for α_2_, *n* = 10 for β_1_, *n* = 9 for FXYD1, and *n* = 6 for FXYD1 phosphorylation (FXYD1/AB_FXYD1).

### Calculations

The plasma K^+^ venous-arterial concentration difference (K_VA_, mM) was calculated by adjusting for changes in the plasma fraction based on haematocrit measurements:

K_VA_ = K_v_ – K_a_ × (1 – Hct_a_) / (1 – Hct_v_), where K_v_ is venous plasma K^+^ concentration, K_a_ is arterial plasma K^+^ concentration, Hct_v_ is venous haematocrit, and Hct_a_ is arterial haematocrit.

The net release of K^+^ from the leg (leg K^+^release, mmol∙min^-1^) was also calculated by adjusting for changes in plasma fraction based on haematocrit measurements:

Net leg K^+^ release = Q̇ × (K_v_ × (1 – Hct_v_) – K_a_ × (1 – Hct_a_)), where Q̇is femoral arterial blood flow in L∙min^-1^.

## Statistics

Data were assessed for normality using graphical model control of histograms. Data that violated normality were log-transformed prior to subsequent analyses. Paired Student’s t-tests were used to test for an effect of time (Pre/Post) on performance (iPPO) for each training condition (CON-leg and BFR-leg). A paired student t-test was used to assess the effect of condition for the change from Pre to Post in performance using the individual changes. For leg blood flow, venous-arterial K^+^ difference, and leg K^+^release, a two-way, repeated-measures (RM) ANOVA was used to test for an effect of time (Pre/Post) and sample no. (Rest, 20 s, 1 min, 3 min, etc.) for each condition. The same test was used to test for an effect of condition for the change from Pre to Post in the same variables using the individual changes. A two-way RM ANOVA was also used to test for an effect of NAC infusion within each condition at Pre and Post, with sample no. and infusion (NAC vs. placebo) as factors, and to assess the effect of NAC over time (Pre vs. Post) within each condition using ∆-infusion values (NAC vs. placebo), with time and sample no. as factors. The same test was used to test for an effect of condition for the effect of NAC using ∆-infusion values (NAC vs. placebo), with condition and sample no. as factors. For protein data, a two-way RM ANOVA was used to test for an effect of time and condition within each fibre type and for the effect of fibre type using time and fibre type as factors. *Post hoc* analyses used the Tukey test. Due to supra-physiological K^+^ values in samples from two individuals caused by late analysis (>5 h) after test termination, *n* = 8 for venous-arterial K^+^ difference and leg K^+^ release. Effect size (*d*) was interpreted based on Cohen’s conventions, where <0.2, 0.2-0.5, >0.5-0.8 and >0.8 were considered as trivial, small, moderate and large effects, respectively (Cohen, 1988). In Results and figures, data are expressed as mean ± SEM, unless otherwise stated. The α-level was set at p ≤ 0.05. Statistical analyses were performed in Sigma Plot (Ver. 11.0, Systat Software, CA), and figures were created using GraphPad Prism 6 (GraphPad Software Inc., California, USA).

## Results

### Effect of BFR on femoral arterial blood flow

In CON-leg, femoral arterial blood flow increased after onset of exercise (20s; *d* = 1.8; p < 0.001; Fig. 2), remained higher during exercise (*d* ≥ 1.7; p < 0.001 vs. rest), and returned to resting level after 40-s of recovery (*d* = 1.8; p = 0.082), after which it remained unchanged (*d* = 0.7-1.6; p ≥ 0.842). In BFR-leg, blood flow increased after 40-s of exercise (*d* = 1.7; p = 0.024 vs. rest), and remained higher during exercise (*d* ≥ 1.8; p ≤ 0.002) and throughout recovery compared to rest (*d* ≥ 1.8; p < 0.001). Blood flow in BFR-leg at rest without (*d* = 0.0; p = 0.984) or with the cuff inflated (*d* = 0.8; p = 0.336) was not different from blood flow at rest in CON-leg. Blood flow in BFR-leg was on average 52 ± 13% lower during exercise (*d* = 1.6; p < 0.001) and 308 ± 113% higher in recovery (*d* = 1.7; p < 0.001) compared to CON-leg.

**Figure 2.**
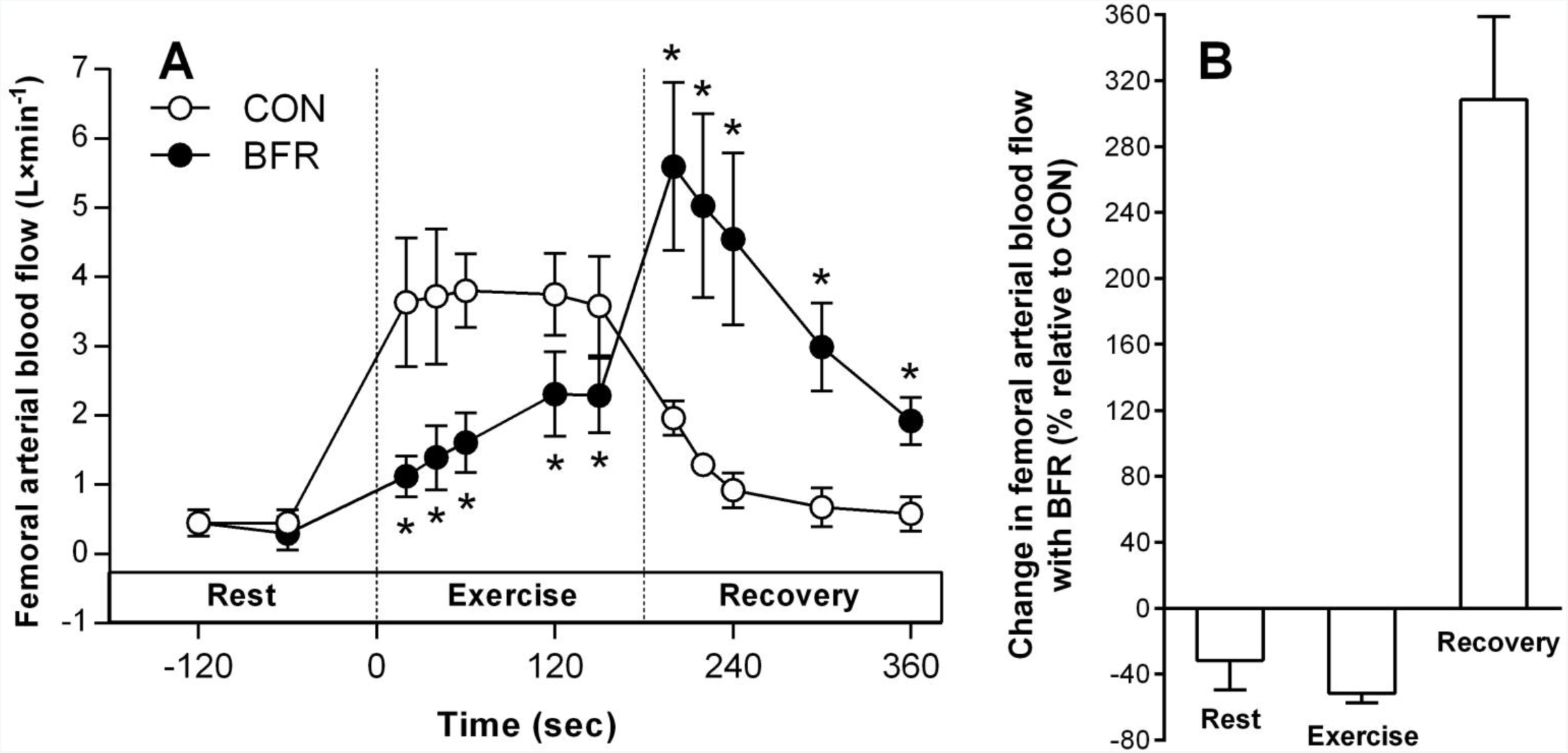
*Acute effects of blood flow restriction on thigh blood flow*. **A)** Femoral arterial blood flow at rest, during exercise, and in recovery without (CON, white symbols) or with (BFR, red symbols) a cuff inflated to ~178 mmHg around the most proximal part of the leg (*n* = 5). *p < 0.001, different from CON. **B)** Percent change in flow induced by blood flow restriction compared to CON.

### Performance

Knee extensor exercise performance (iPPO) increased with the training period by 11 ± 12% in CON-leg (p = 0.043; *d* = 0.5) and by 23 ± 15% in BFR-leg (p < 0.001; *d =* 0.9), with the increase being 12 ± 10% greater in BFR-leg compared to CON-leg (*d =* 0.9; p = 0.034; Fig. 3A). Changes in performance of each leg, represented by the same symbol for each individual, are shown in Fig. 3B.

**Figure 3.**
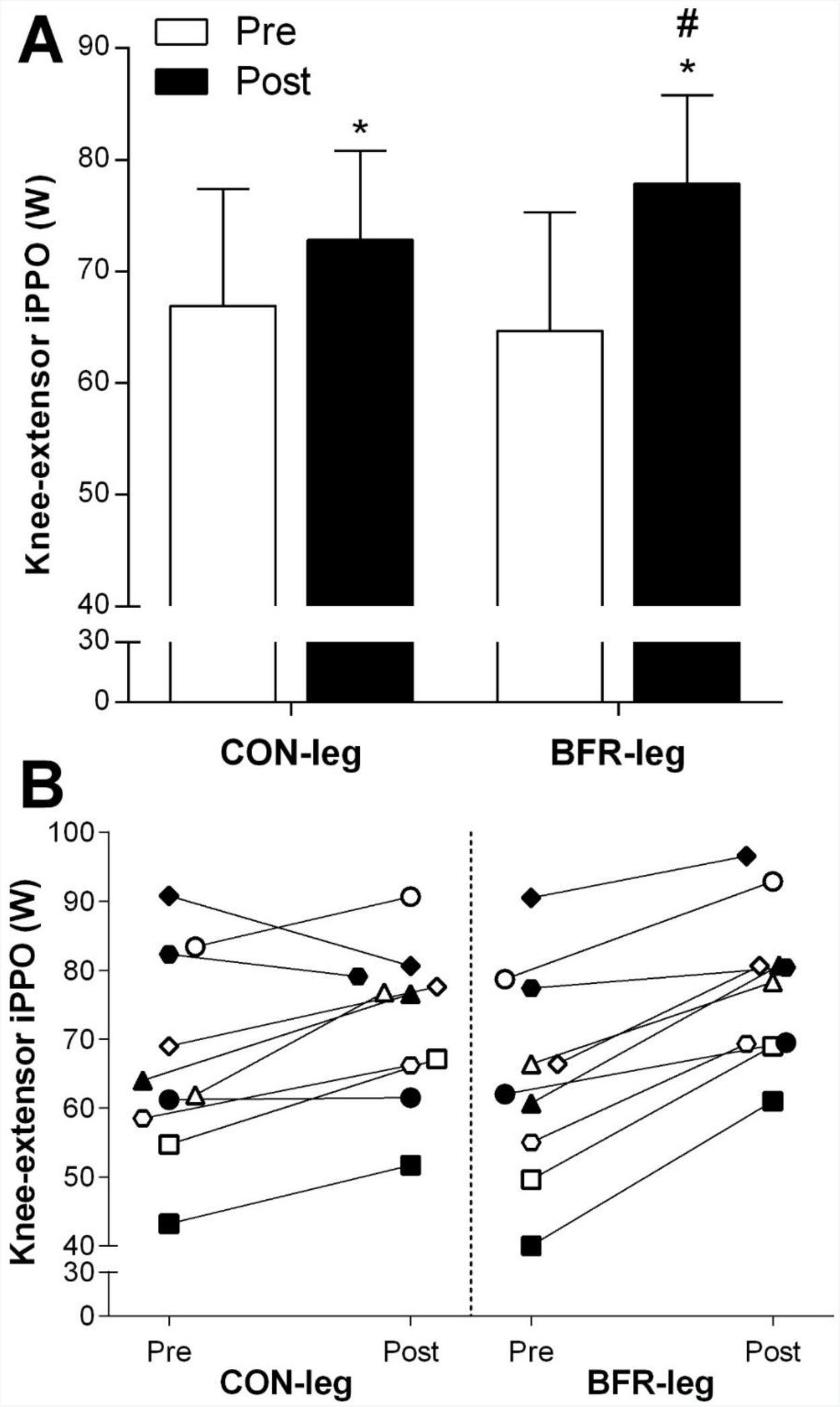
*Effect of training with blood flow restriction on performance during single-leg exercise*. **A)** Knee-extensor incremental peak aerobic power output (iPPO) for the control (CON; *n* = 10) and BFR-trained leg (BFR; *n* = 10) before (Pre, white bars) and after (Post, black bars) training. **B)** Individual changes in iPPO for CON and BFR, with each symbol representing the same individual. *p < 0.05, different from Pre; # p < 0.05, different from CON for the change from Pre to Post. Data are expressed as either mean ± 95 % confidence intervals (**A**) or as absolute values (**B**).

### Femoral arterial blood flow

Blood flow at rest before, during, and in recovery from Ex1 did not change with the training period in CON-leg (*d* = 0.1; p = 0.226; Fig. 4A), but was higher during exercise in BFR-leg after the training period compared with before (20s, 1 min, 8 min: *d* = 0.4-0.6; p ≤ 0.034; Fig. 4B). After the training period, blood flow in BFR-leg was higher than in CON-leg 20-s (*d* = 0.9; p = 0.035) and 8 min into Ex1 (*d* = 0.8; p < 0.001). During Ex2, blood flow was higher at +20-s in CON-leg after the training period compared to before (*d* = 0.4; p < 0.001; Fig. 4C) and in BFR-leg at 1 min (*d* = 0.5; p = 0.029; Fig. 4D), at which point blood flow was higher in BFR-leg compared to CON-leg (*d* = 1.1; p = 0.029).

**Figure 4.**
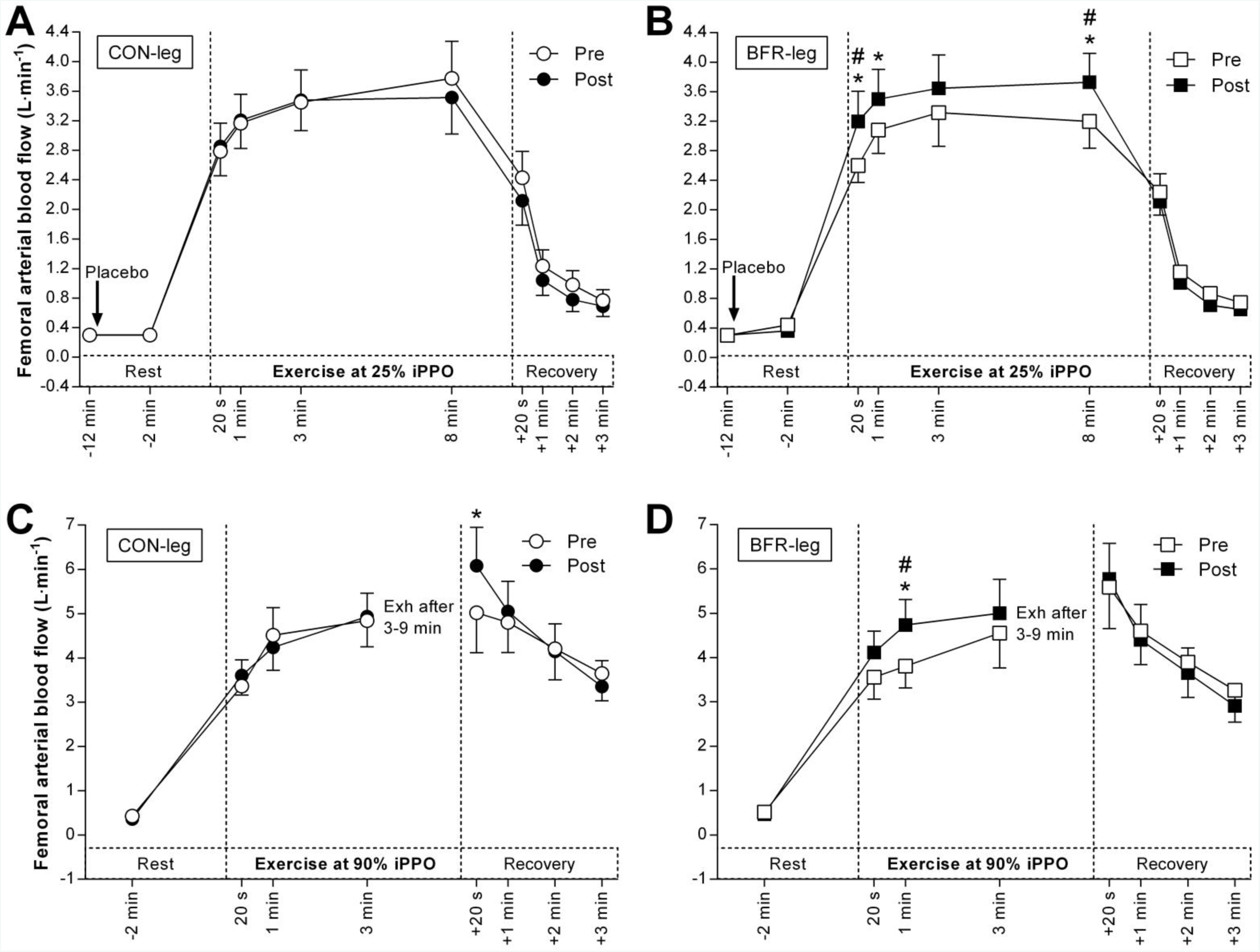
*Effect of training with blood flow restriction on thigh blood flow during exercise*. **A)** Blood flow in the leg that trained without blood flow restriction (CON-leg, circles) during exercise at 25% incremental peak power output (iPPO) before (Pre, white symbols) and after (Post, black symbols) training. **B)** Blood flow in the leg that trained with blood flow restriction (BFR-leg, squares) during exercise at the same intensity. **C)** Blood flow in CON-leg during exercise at 90% iPPO to exhaustion (Exh). **D)** Blood flow in BFR-leg during exercise at the same intensity. *n* = 10. The arrow indicates the start of infusion. *p < 0.05, different from Pre; #p < 0.05, greater change from Pre to Post compared to CON-leg. Data are expressed as mean ± SEM.

### Venous-arterial K^+^ difference (K_VA_)

K_VA_ did not change with the training period in CON-leg, neither during Ex1 (*d* = 0.0; p = 0.885; Fig. 5A) nor Ex2 (p = 0.965, *d* = 0.1; Fig. 5C). In contrast, K_VA_ was lower in BFR-leg after the training period, both during Ex1 (20s: *d* = 0.6; p = 0.037; 3 min: *d* = 0.6; p = 0.065; Fig. 5B) and Ex2 (1 min: *d* = 0.9; p = 0.050; 3 min: *d* = 1.0; p < 0.001), and in recovery from Ex2 (*d* = 0.8-1.3; p ≤ 0.029; Fig. 5D). The decline in K_VA_ with training in BFR-leg at 3 min of Ex2 (*d* = 0.8; p = 0.014) and 3 min after Ex2 (p < 0.001, *d* = 0.9) was larger than the corresponding change in CON-leg during the training period (Fig. 5D).

**Figure 5.**
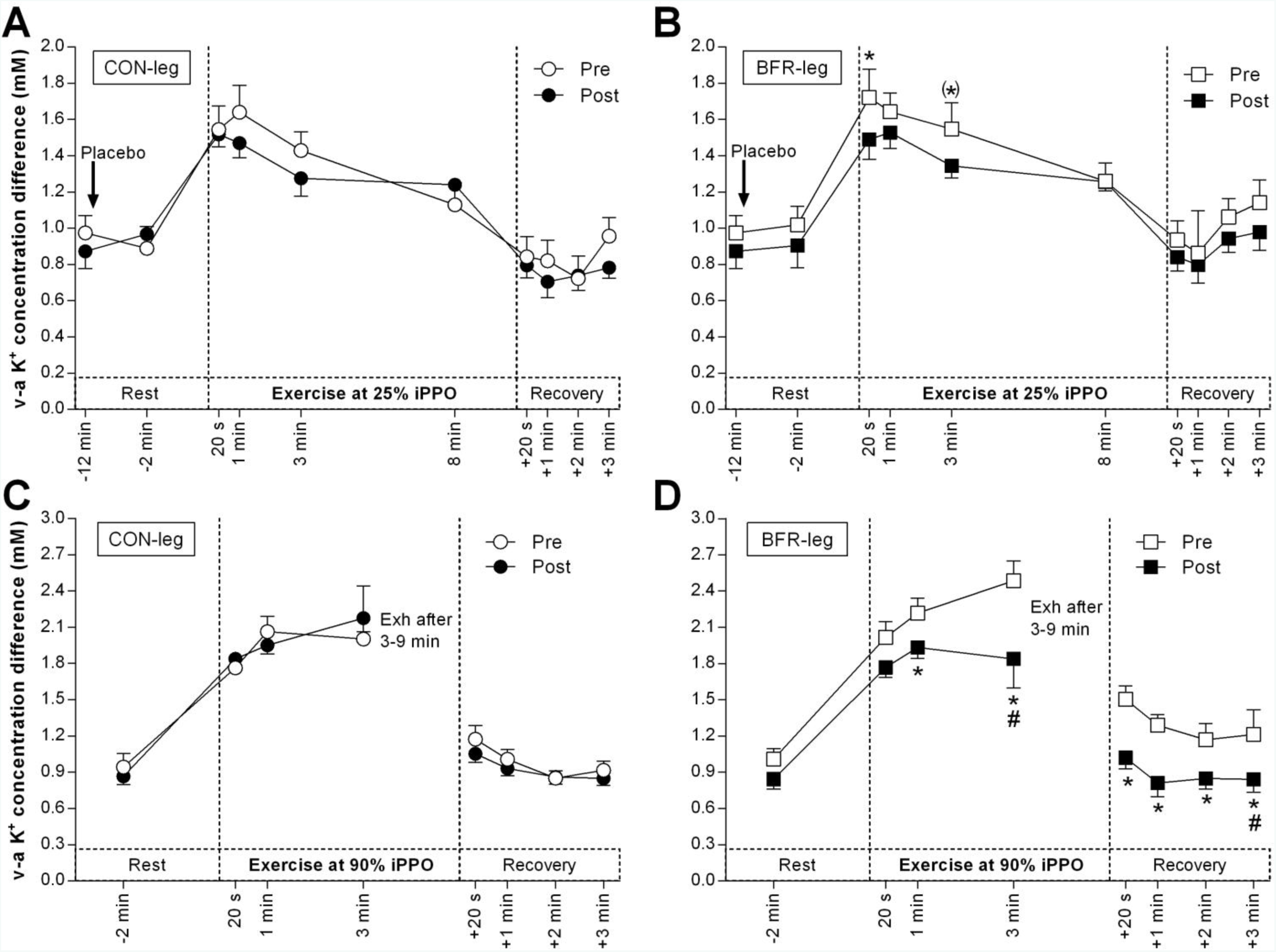
*Effect of training with blood flow restriction on venous-arterial (v-a) K^+^ difference during exercise*. **A)** v-a K^+^ difference in the leg that trained without blood flow restriction (CON-leg, circles) during exercise at 25% incremental peak power output (iPPO) before (Pre, white symbols) and after (Post, black symbols) training. **B)** v-a K^+^ difference in the leg that trained with blood flow restriction (BFR-leg, squares) during exercise at the same intensity. **C)** v-a K^+^ difference in CON-leg during exercise at 90% iPPO to exhaustion (Exh). **D)** v-a K^+^ difference in BFR-leg during exercise at the same intensity. *n* = 8. The arrow indicates the start of the infusion. *p < 0.05, different from Pre; (*) p = 0.065, different from Pre; #p < 0.05, greater change from Pre to Post compared to CON-leg. Data are expressed as mean ± SEM.

### Thigh K^+^ release

In Ex1, net K^+^ release did not change with the training period in both legs (*d* ≤ 0.6; p ≥ 0.600; Fig. 6), with no differences between legs (*d* ≤ 0.8; p ≥ 0.142). In Ex2, net K^+^ release did not change with the training period in CON-leg (*d* = 0.2; p = 0.525), but was reduced at 3 min in BFR-leg after the training period (*d* = 0.9; p = 0.004), with the decrease being larger than in CON-leg at this time point (*d* = 0.9; p = 0.005).

**Figure 6.**
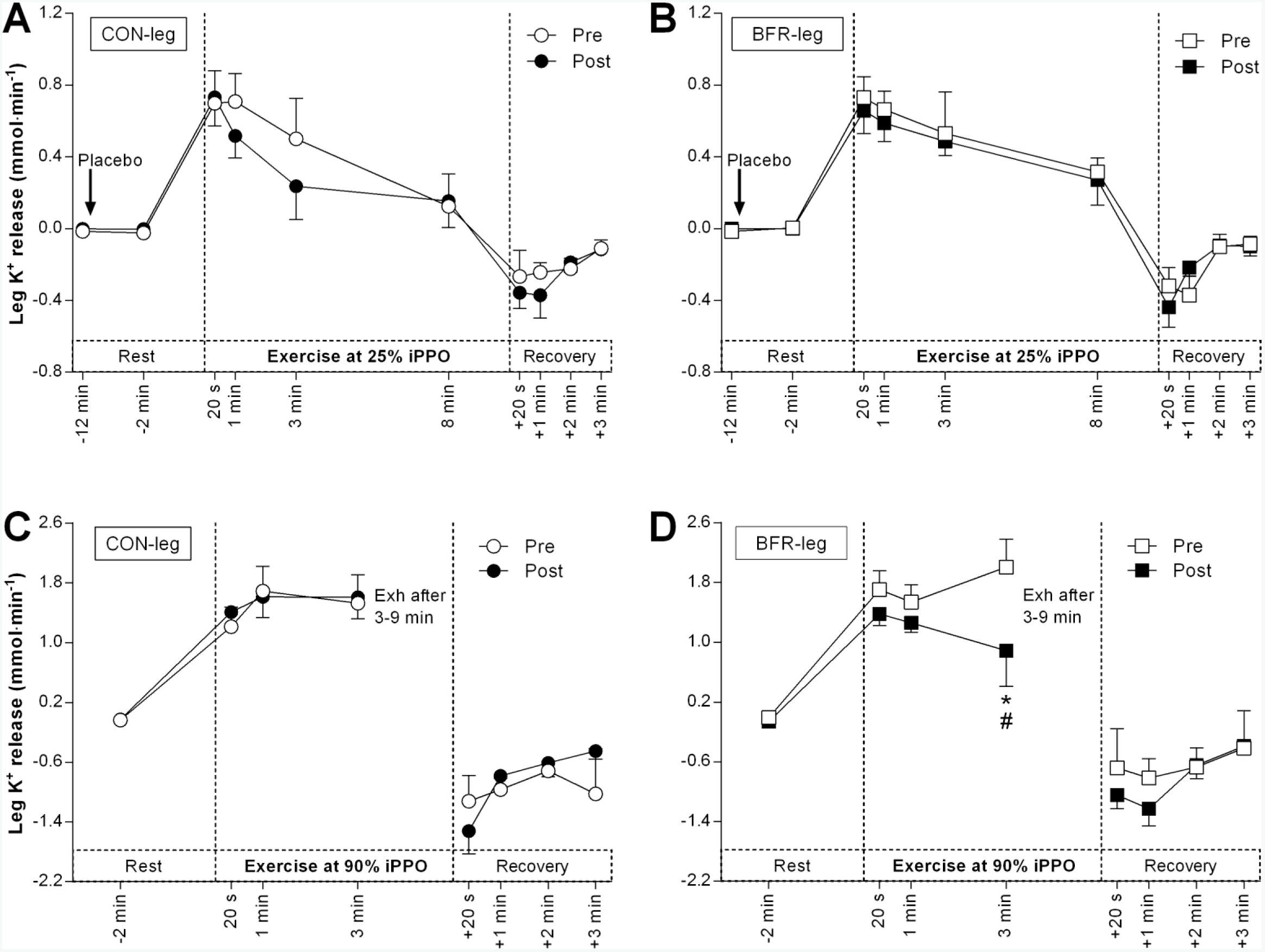
*Effect of training with blood flow restriction on net thigh K^+^ release during exercise*. **A)** K^+^ release from the leg that trained without blood flow restriction (CON-leg, circles) during exercise at 25% incremental peak power output (iPPO) before (Pre, white symbols) and after (Post, black symbols) training. **B)** K^+^ release from the leg that trained with blood flow restriction (BFR-leg, squares) during exercise at the same intensity. **C)** K^+^ release from the CON-leg during exercise at 90% iPPO to exhaustion (Exh). **D)** K^+^ release from BFR-leg during exercise at the same intensity. *n* = 8. The arrow indicates the start of the infusion of saline. *p < 0.05, different from Pre; #p < 0.05, greater change from Pre to Post compared to CON-leg. Data are expressed as mean ± SEM.

### Effects of NAC infusion on femoral arterial blood flow during exercise at 25% iPPO

Before the training period, blood flow in CON-leg was lower with NAC compared to PLA infusion during Ex1 (8 min: p = 0.004, *d* = 0.2; Fig. 7A), and in recovery from Ex1 (+20s: *d* = 0.4; p = 0.001; +1 min: *d* = 0.5; p = 0.005; +2 min: *d* = 0.5; p = 0.012). In BFR-leg, blood flow was higher with NAC compared to PLA infusion during Ex1 (20s: *d* = 0.4; p = 0.040; 8 min: *d* = 0.4; p = 0.004; Fig. 7B), with the effect of NAC being greater in BFR-leg than in CON-leg after 8 min of Ex1 (*d* = 1.1; p < 0.001).

**Figure 7.**
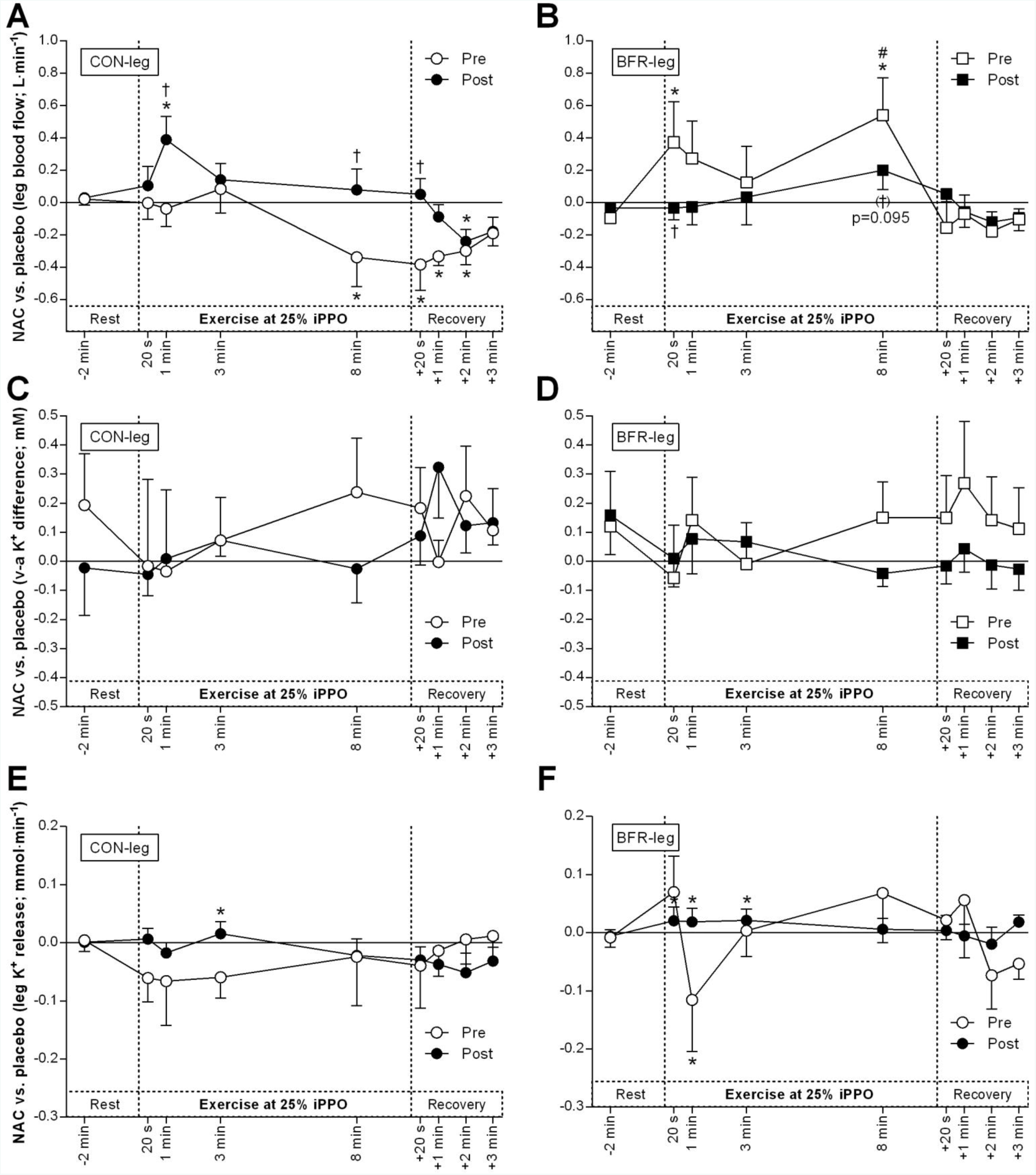
*Effect of antioxidant infusion on thigh blood flow, venous-arterial (v-a) K^+^ difference, and net K^+^ release during exercise at 25% incremental peak aerobic power output before and after training with blood flow restriction*. Difference between N-acetylcysteine (NAC) and saline (placebo) infusion in femoral arterial blood flow (**A** + **B**; *n* = 10), v-a K^+^ difference (**C** + **D**; *n* = 8), and thigh K^+^ release (**E** + **F**; *n* = 8) before (Pre, white) and after (Post, black) cycling without (CON-leg, circles) or with blood flow restriction (BFR-leg, squares). *p < 0.05, different from placebo; #p < 0.05, different from CON-leg; † p < 0.05, different from Pre. Data are expressed as mean ± SEM.

After the training period, blood flow in CON-leg was higher with NAC compared to PLA infusion during Ex1 (1 min: *d* = 0.3; p < 0.001) and lower in recovery from Ex1 (+2 min: *d* = 0.5; p = 0.020; Fig. 7A). The effect of NAC on blood flow in CON-leg was greater after than before the training period during Ex1 (1 min: p = 0.003; 8 min: p = 0.003), and in recovery from Ex1 (+20s: p = 0.002; Fig. 7A). After the training period, NAC infusion did not change blood flow in BFR-leg compared to PLA during Ex1 (*d* = 0.3; p ≥ 0.099; Fig. 7B). Further, the effect of NAC on blood flow in BFR-leg was lower after relative to before the training period after 20s of Ex1 (p = 0.047), and tended to be lower after 8 min of Ex1 (p = 0.095; Fig. 7B). However, the effect of NAC on blood flow was not different between the legs during Ex1 after the training period (*d* = 0.1; p = 0.442).

### Effects of NAC infusion on femoral arterial blood flow during exercise at 90% iPPO

Before the training period, blood flow in CON-leg did not change with NAC compared to PLA infusion during Ex2 (*d* = 0.0; p = 0.988; Fig. 8A). Blood flow in BFR-leg was higher with NAC compared to PLA infusion during Ex2 (20s: *d* = 0.4; p = 0.046; 1 min: *d* = 0.6; p = 0.005; 3 min: *d* = 0.3; p = 0.058), and in recovery from Ex2 (1 min: *d* = 0.3; p = 0.025; Fig. 8B). The effect of NAC on blood flow was greater in BFR-leg than in CON-leg after 1 min of Ex2 before the training period (*d* = 0.5; p = 0.006; Fig. 8B).

**Figure 8.**
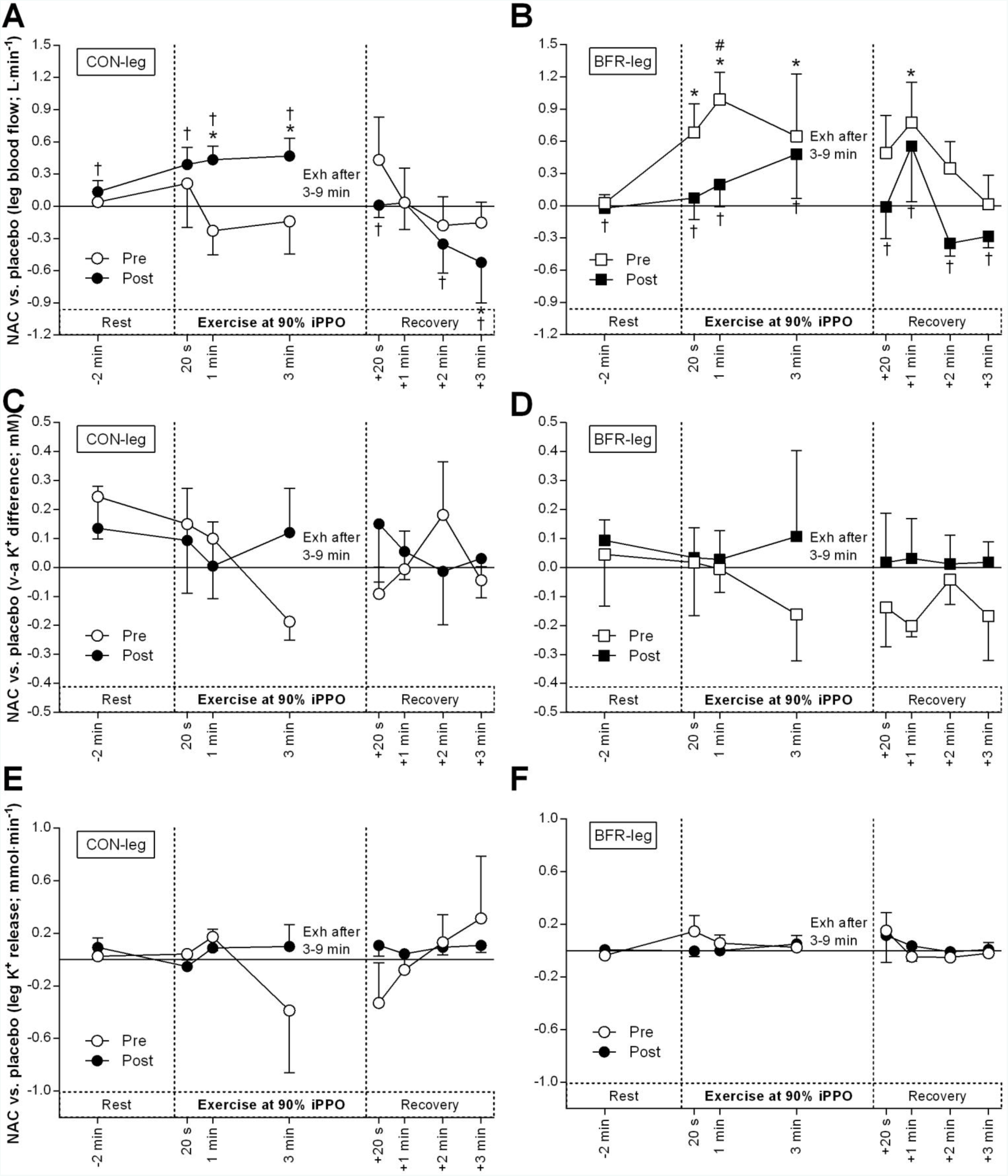
*Effect of antioxidant infusion on thigh blood flow, venous-arterial (v-a) K^+^ difference, and net K^+^ release during exercise at 90% incremental peak aerobic power output to exhaustion (Exh) before and after cycling with blood flow restriction*. Difference between N-acetylcysteine (NAC) and saline (placebo) infusion in femoral arterial blood flow (**A** + **B**; *n* = 10), v-a K^+^ difference (**C** + **D**; *n* = 8), and K^+^ release (**E** + **F**; *n* = 8) before (Pre, white) and after (Post, black) cycling without (CON-leg, circles) or with blood flow restriction (BFR-leg, squares). *p < 0.05, different from placebo; #p < 0.05, different from CON-leg; † p < 0.05, different from Pre. Data are expressed as mean ± SEM.

After the training period, blood flow in CON-leg was higher with NAC compared to PLA infusion during Ex2 (1 min: *d* = 0.3; p = 0.048; 3 min: *d* = 0.2; p = 0.033) and in recovery from Ex2 (3 min: *d* = 0.3; p = 0.018; Fig. 8A). The effect of NAC on blood flow in CON-leg was greater after compared to before the training period at rest before (p < 0.001), during (20s, 1 min, and 3 min: p < 0.001), and in recovery from Ex2 (+20s, +1 min, +2 min, and +3 min: p < 0.001). After the training period, blood flow in BFR-leg did not change with NAC compared to PLA infusion during Ex2 (*d* = 0.0; p = 0.463; Fig. 8B). The effect of NAC on blood flow in BFR-leg was lower after compared to before the training period at all time points during Ex2 (p ≤ 0.002). However, the effect of NAC on blood flow was not different between the legs during Ex2 after the training period (*d* = 0.0; p = 0.967).

### Effects of NAC infusion on venous-arterial K^+^ difference, K_VA_

Before the training period, K_VA_ in CON-leg did not change with NAC compared to PLA infusion during both Ex1 (*d* = 0.1; p = 0.334; Fig. 7C) and Ex2 (*d* = 0.2; p = 0.627; Fig. 8C). Similarly, K_VA_ in BFR-leg remained unaffected by NAC compared to PLA infusion during Ex1 (*d* = 0.1; p = 0.628; Fig. 7D) and Ex2 (*d* = 0.2; p = 0.732; Fig. 8D). Before the training period, the effect of NAC was not different between the legs both during Ex1 (*d* = 0.2; p = 0.190) and during Ex2 (*d* = 0.1; p = 0.680).

After the training period, K_VA_ in CON-leg did not change with NAC compared to PLA infusion during both Ex1 (*d* = 0.2; p = 0.355; Fig. 7C) and Ex2 (*d* = 0.1; p = 0.343; Fig. 8C). The effect of NAC on K_VA_ in CON-leg was not different after compared to before the training period during both Ex1 (p = 0.387) and Ex2 (p = 0.659). Similarly, K_VA_ in BFR-leg did not change with NAC compared to PLA infusion during Ex1 (*d* = 0.1; p = 0.207; Fig. 7D) and Ex2 (*d* = 0.1; p = 0.252; Fig. 8D) after the training period, and the effect of NAC on K_VA_ in BFR-leg was not different after compared to before training during both Ex1 (p = 0.824) and Ex2 (p = 0.194). After the training period, the effect of NAC was not different between the legs both during Ex1 (*d* = 0.2; p = 0.348) and during Ex2 (*d* = 0.2; p = 0.233).

### Effects of NAC infusion on thigh K^+^ release

Before the training period, net K^+^ release in CON-leg did not change with NAC compared to PLA infusion during both Ex1 (d ≤ 1.0; p = 0.635; Fig. 7E) and Ex2 (*d* = 0.1; p = 0.914; Fig. 8E), although the large effect sizes during Ex1 should be noted. In BFR-leg, net K^+^ release decreased with NAC compared to PLA infusion after 1 min of Ex1 (*d* = 0.7; p = 0.041; Fig. 7F), whereas net K^+^ release during Ex2 was unaffected by NAC (*d* = 0.1; p = 0.812; Fig. 8F). Before the training period, no differences between legs were detected for the effect of NAC during either Ex1 (*d* = 0.2; p = 0.642) or Ex2 (*d* = 0.4; p = 0.164).

After the training period, net K^+^ release in CON-leg increased with NAC compared to PLA infusion after 3 min of Ex1 (p = 0.011, *d* = 0.6; Fig. 7E), but remained unaffected by NAC during Ex2 (*d* = 0.1; p = 0.640; Fig. 8E). In CON-leg, the effect of NAC on net K^+^ release was not different after compared to before the training period during either Ex1 (p = 0.300) or Ex2 (p = 0.615). After the training period, net K^+^ release in BFR-leg increased with NAC compared to PLA infusion during Ex1 (20s: *d* = 0.5; p = 0.056; 1 min: *d* = 1.0; p < 0.001; 3 min: *d* = 0.6; p = 0.054; Fig. 7F), but did not change with NAC during Ex2 (*d* = 0.2; p = 0.812; Fig. 8F). In BFR-leg, the effect of NAC on net K^+^ release was not different after compared to before the training period during either Ex1 (p = 0.420; Fig. 7F) or Ex2 (p = 0.313; Fig. 8F). No differences were detected between the legs for an effect of NAC on net K^+^ release after the training period during either Ex1 (*d* = 0.4; p = 0.127) or Ex2 (*d* = 0.3; p = 0.112).

### Na^+^,K^+^-ATPase α-isoform abundance in type I and II muscle fibres

Representative blots for all proteins are shown in Fig. 9K.

In CON-leg, α_1_ abundance did not change with training in either type-I (30%, *d* = 0.5; p = 0.387; Fig. 9A) or type-II fibres (7%, *d* = 0.2; p = 0.508; Fig. 9B). In BFR-leg, α_1_ abundance tended to increase in type-I (46%, *d* = 0.7; p = 0.075; Fig. 9A), but not in type-II fibres (26%, *d* = 0.5; p = 0.508; Fig. 9B). In type-I fibres, no difference was detected between legs for α_1_ abundance either before (1%) or after (11%) the training period (*d* ≤ 0.2; p = 0.662). However, in type-II fibres, α_1_ abundance was higher in BFR-leg compared to CON-leg after the training period (51%, *d* = 0.7; p = 0.002). An effect of fibre type was evident for α_1_ abundance in CON-leg, with a higher abundance in type-I than in type-II fibres (29%, *d* = 0.7; p = 0.047), whereas no effect of fibre type was observed in BFR-leg (5%, *d* = 0.2; p = 0.505).

**Figure 9.**
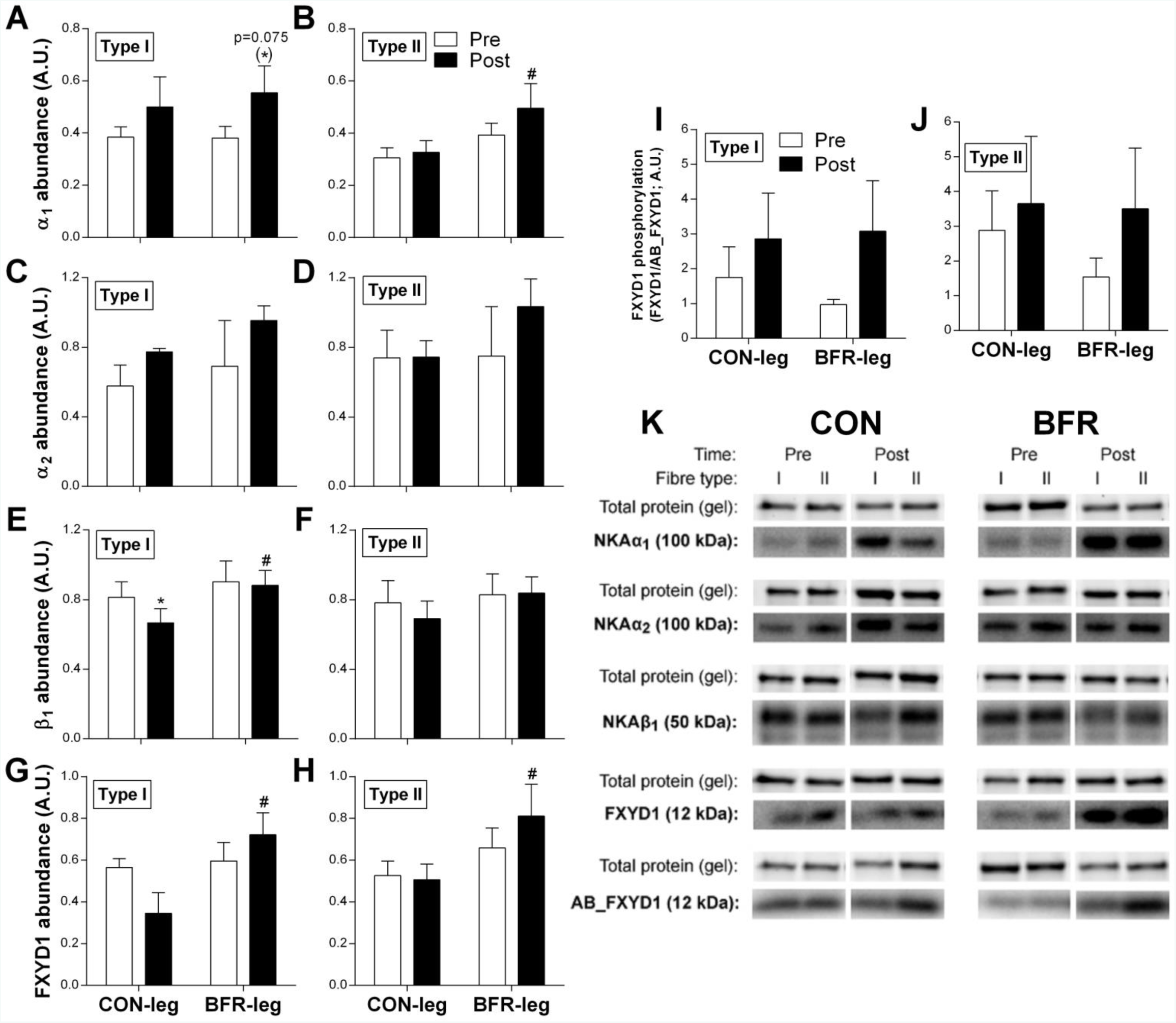
*Effect training with blood flow restriction on Na^+^,K^+^-ATPase-isoform abundance in type I and II muscle fibres of men.* α_1_ (**A** + **B**; *n* = 9), α_2_ (**C** + **D**; *n* = 3), β_1_ (**E** + **F**; *n* = 10) and FXYD1 abundance (**G** + **H**; *n* = 9), and FXYD1 phosphorylation (FXYD1/AB_FXYD1; **I** + **J**; *n = 6*) was determined in type I and II muscle fibres from a leg cycling without (CON-leg) or with blood flow restriction (BFR) for six weeks both before (Pre, white bars) and after (Post, black bars) training. *p ≤ 0.05, different from Pre. #p ≤ 0.05, different from CON at Post. †p ≤ 0,05, different from type-II fibres at Post Data are expressed as mean ± SEM. Representative western blots for measured proteins are shown in **K**. Total protein was determined as total protein content in each lane on the stain-free gel image, represented here as the Actin-band.

In CON-leg, α_2_ abundance did not change with training in either type-I (35%, *d* = 1.2; p = 0.411; Fig. 9C) or type-II fibres (1%, *d* = 0.02; p = 0.690; Fig. 9D). Similarly, in BFR-leg, α_2_ abundance did not change in either type-I (38%, *d* = 0.8; p = 0.411; Fig. 9C) or type-II fibres (38%, *d* = 0.7; p = 0.690; Fig. 9D), despite the large effect sizes and magnitude of increases. No difference was evident between legs for α_2_ abundance in either type-I (*d* ≤ 1.3, p ≥ 0.301) or type-II fibres (*d* ≤ 1.1, p = 0.137). In addition, no effect of fibre type for α_2_ abundance was detected in either leg (*d* ≤ 0.7; p ≥ 0.370).

### Na^+^,K^+^-ATPase β_1_ and FXYD1 abundance in type I and II muscle fibres

In CON-leg, β_1_ abundance decreased with training in type-I (–18%, *d* = 0.5; p = 0.022; Fig. 9E), but did not change in type-II fibres (–12%, *d* = 0.3; p = 0.542; Fig. 9F). In BFR-leg, β_1_ abundance did not change in either type-I (–2%, *d* = 0.1; p = 0.827; Fig. 9E) or type-II fibres (1%, *d* = 0.03; p = 0.542; Fig. 9F). After the training period, β_1_ abundance in type-I fibres was higher in BFR-leg than in CON-leg (33%, *d* = 0.8; p = 0.019), whereas no difference between legs was detected in type-II fibres (*d* ≤ 0.5; p = 0.084). In addition, no effect of fibre type was evident for β_1_ abundance in both legs (*d* ≤ 0.2; p ≥ 0.359).

In CON-leg, FXYD1 abundance did not change with training in type-I (–39%, *d* = 0.9; p = 0.603; Fig. 9G) or type-II fibres (–4%, *d* = 0.1; p = 0.603; Fig. 9H), despite the large effect size and mean decrease in type-I fibres. In BFR-leg, FXYD1 abundance did not change with training in type-I (21%, *d* = 0.5; p = 0.603; Fig. 9G) or type-II fibres (24%, *d* = 0.4; p = 0.761; Fig. 9H), despite the substantial mean gains and moderate effect sizes. After the training period, FXYD1 abundance was higher in BFR-leg than in CON-leg in both type-I (108%, *d* = 1.1; p = 0.025) and type-II fibres (60%, *d* = 0.8; p = 0.035). An effect of fibre type was evident in BFR-leg, with FXYD1 abundance being higher in type-II compared to type-I fibres (12%, *d* = 0.3; p = 0.017), but no effect of fibre type was observed in CON-leg (–13% vs. type-I, *d* ≤ 0.6; p = 0.233).

### FXYD1 phosphorylation in type I and II muscle fibres

In CON-leg, FXYD1 phosphorylation, as expressed relative to total FXYD1 abundance (FXYD1/AB_FXYD1), did not change with training in either type-I (63%, *d* = 0.5; p = 0.166; Fig. 9I) or type-II fibres (27%, *d* = 0.2; p = 0.402; Fig. 9J). Similarly, in BFR-leg, FXYD1 phosphorylation did not change with training in either type-I (217%, *d* = 0.8; p = 0.166; Fig. 9I) or type-II fibres (127%, *d* = 0.6; p = 0.402; Fig. 9J), despite the large mean increases and effect sizes. No difference between legs was detected for FXYD1 phosphorylation in either type-I (*d* ≤ 0.5; p = 0.586) or type-II fibres (*d* ≤ 0.6; p = 0.502). An effect of fibre type for FXYD1 phosphorylation was observed in BFR-leg, with a higher abundance in type-II than in type-I fibres (25%, *d* = 0.6; p = 0.017), whereas no effect of fibre type was evident in CON-leg (47%, *d* = 0.5; p = 0.250).

## Discussion

The novel findings of the present study were that six weeks of interval cycling with reduced blood flow to exercising muscles elicited greater improvements in single-leg exercise performance compared to training without blood flow restriction (BFR). Furthermore, training with BFR did, in contrast to control, reduce muscle K^+^ release during near-maximal exercise. Before training, antioxidant infusion depressed thigh K^+^ release in the initial phase of moderate-intensity exercise, whereas after training, the antioxidant effect was blunted in the BFR-trained leg. In addition, the reduced net thigh K^+^ release after training with BFR was associated with higher abundance of some Na^+^,K^+^-ATPase isoforms in type-I (β_1_: 33%, FXYD1: 108%) and type-II (α_1_: 51%, FXYD1: 60%) muscle fibres.

### Improvements in performance and thigh K^+^ handling by training with BFR

Despite performing the same amount of work, the leg that trained with BFR improved exercise performance substantially more (12%) than the leg training without BFR. Thus, interval cycling with BFR incorporating rest periods with no circulatory restriction is an effective strategy to augment training-induced improvements in exercise tolerance. Similar enhancing effects of BFR have been observed in studies that used other occlusion training modalities. By use of local supra-atmospheric pressure (+50 mmHg) to reduce (~16%) blood flow to exercising muscles, Sundberg *et al.* (1993) found an 11% greater increase in time to exhaustion during single-leg exercise for a leg training with compared to a leg training without reduced blood flow (25 vs. 14 %; 45 min /session, 4 days/week for 4 weeks). In agreement, fifteen minutes of bilateral cycling (40% VO_2max_) with BFR induced by inflation of a cuff around the legs (pressure: 160-220 mmHg; 3 days/week for 8 weeks) increased cycling time to exhaustion by 15%, whereas no performance change was detected in a control group training without BFR, despite this group had a greater (66%) training volume (Abe *et al.*, 2010).

Enhancements in performance of the BFR-trained leg was associated with reduced net thigh K^+^ release rate from exercising muscles during intense exercise. The lower rate of thigh K^+^ release may be explained by training-induced adaptations in Na^+^,K^+^-ATPase α_1_, β_1_ and FXYD1 abundances in type I and α_1_ and FXYD1 in type II muscle fibres, which may increase the potential for assembly of a greater number of Na^+^,K^+^-ATPase units in the sarcolemma and T-tubuli during exercise, thereby increasing the capacity for K^+^ re-uptake (Clausen, 2003). Furthermore, FXYD1 phosphorylation tended to increase in both type I (217%, *d* = 0.8; p = 0.166) and II fibres (127%, *d* = 0.6; p = 0.402) in the BFR-trained leg. Given that phosphorylation of FXYD1 increases Na^+^ affinity and activity of the Na^+^,K^+^-ATPase (Bibert *et al.*, 2008; Ingwersen *et al.*, 2011), a higher FXYD1 phosphorylation may have promoted Na^+^,K^+^-ATPase activation, and hence contributed to the reduced rate of K^+^ release from the BFR-trained leg observed during intense exercise. However, the small sample size (*n* = 6) for the measurement of FXYD1 phosphorylation likely precluded the change from being significant and the interpretation should be regarded with caution.

Another adaptation that may have contributed to the reduced net release of K^+^ from the muscles in the BFR-trained leg is the faster rise in blood flow to the exercising muscles at onset of exercise. This is supported by findings that microvascular permeability for K^+^ is positively related to blood flow (Friedman & DeRose, 1982; Kajimura *et al.*, 1998). Thus, higher flow would lead to faster and/or greater removal of K^+^ from the interstitial space, thereby facilitating re-establishment of the transmembrane K^+^ gradient. Studies have shown that BFR-training increases muscle capillary density and tissue perfusion (Sundberg, 1994; Kacin & Strazar, 2011), and induces angiogenic factors (Larkin *et al.*, 2012). As such, a greater surface area for diffusion of K^+^ from the muscle interstitium to the blood stream may be expected after BFR-training. This is supported by the substantial increase (>3 fold) in femoral arterial blood flow after cuff deflation (Fig. 2) and resultant high shear stress. However, we observed lower rates of venous and arterial blood K^+^ accumulation in the BFR-trained leg (*Suppl. table 1 and 2*), indicating K^+^ re-uptake was a significant contributor to the attenuated net K^+^ release from exercising muscles after BFR-training.

### Role of ROS in regulating muscle K^+^ handling in humans and effects of BFR-training

Before training, intravenous infusion of the antioxidant NAC depressed net K^+^ release during moderate-intensity exercise in the BFR-leg. A similar, but non-significant trend was observed in the control leg (large effect size: ~1), which possibly was related to a large inter-subject variation in the response to NAC (Paschalis *et al.*, 2017). The observation that antioxidant infusion reduces thigh K^+^ release during exercise extends previous findings of decreased systemic (arterialised-venous) K^+^ concentrations during moderate-intensity cycling in response to NAC (Medved *et al.*, 2004; McKenna *et al.*, 2006). Together, these data indicate that ROS alters K^+^ handling in exercising muscles in humans.

After the six weeks of training with BFR, net thigh K^+^ release was reduced concomitant with a blunted effect of antioxidant infusion on net K^+^ release. This suggests the improved K^+^ handling in the BFR-trained leg may have been caused, in part, by increasing the capacity to scavenge ROS. Consistent with the reduced effect of NAC on net K^+^ release in the BFR-trained leg, both a better K^+^ handling by exercising muscles (Nielsen *et al.*, 2004) and increased muscle content and/or activity of antioxidants have been observed after high-intensity training in humans (Hellsten *et al.*, 1996; Leeuwenburgh *et al.*, 1997). An association has also been observed between Na^+^,K^+^-ATPase function and degree of glutathionylation (oxidative stress) of the β-subunit following intense cycling to exhaustion in endurance-trained men (Juel *et al.*, 2015).

The training-induced increases in FXYD1 abundance may also have contributed to the reduced effect of NAC after the training intervention, since FXYD1 overexpression has been shown to protect against ROS-induced Na^+^,K^+^-ATPase inhibition by de-glutathionylation of the β-subunit in both myocytes (Liu et al., 2013) and *Xenopus* oocytes (Bibert *et al.*, 2011). In addition, susceptibility of the Na^+^,K^+^-ATPase to oxidative inhibition by glutathionylation was depressed with increasing K^+^ concentration *in vitro* (Liu *et al.*, 2012). Thus, when K^+^ homeostasis is severely perturbed by raising extracellular K^+^ levels, such as during strenuous exercise (Nielsen *et al.*, 2004; Gunnarsson *et al.*, 2013), the effect of ROS on Na^+^,K^+^-ATPase function becomes less potent. This may explain the absence of effect of NAC on net thigh K^+^ release at the high exercise intensity in the present study. Further, it should be considered that altered function of other K^+^ regulatory systems, such as the Na^+^–K^+^–2CL^-^ exchanger (NKCC1), inward rectifying K^+^ channel (K_IR_2.1) and K_ATP_ channel (K_IR_6.2), may also have contributed to blunt the effect of NAC and improve K^+^ handling after training with BFR, since these systems are also redox sensitive (Kourie, 1998) and their expression altered by training in humans (Hostrup & Bangsbo, 2017).

### Training-induced adaptations in Na^+^,K^+^-ATPase isoforms in type I and II fibres

Recent work with different types of training, including sprint-interval (Wyckelsma *et al.*, 2015; Christiansen *et al.*, 2017b), resistance (Perry *et al.*, 2016), and interval endurance (Wyckelsma *et al.*, 2017), have shown that the abundance of Na^+^,K^+^-ATPase isoforms are altered in a fibre-type specific manner by training in human skeletal muscle. However, these measurements have not been linked to K^+^ transport, which was done in the present study. The divergent, fibre type-specific adaptations elicited by BFR-training compared to training without BFR probably reflect altered modulation of the cellular stressors and signalling pathways that regulate Na^+^,K^+^-ATPase-isoform expression. For example, BFR was shown to promote exercise-induced downstream AMPK signalling and oxidative stress, which was associated with a greater mRNA response of FXYD1 in trained men (Christiansen *et al.*, 2018), indicating ROS and AMPK could be important factors for mediating increases in FXYD1 expression after training with BFR.

### Summary

Six weeks of interval cycling with BFR elicited greater gains in single-leg exercise performance compared to worked-matched training without BFR, which was associated with reduced rate of K^+^ release from the thigh during intense exercise. The enhanced K^+^ handling after BFR-training may be related to improved protection against oxidative damage, as indicated by an attenuated effect of antioxidant treatment on net thigh K^+^ efflux, and to higher abundance of Na^+^,K^+^-ATPase isoforms in type I (α_1_, β_1_, FXYD1) and II muscle fibres (α_1_, FXYD1). Furthermore, increases in blood flow to exercising muscles and FXYD1 phosphorylation might also have contributed to the effect of training with BFR on K^+^ handling. In addition, the current data indicate that ROS alters K^+^ handling in exercising muscles in humans. Based on the present findings, a visual summary of the proposed factors underlying the improved skeletal muscle K^+^ handling by BFR-training is shown in Fig. 10.

**Figure 10.**
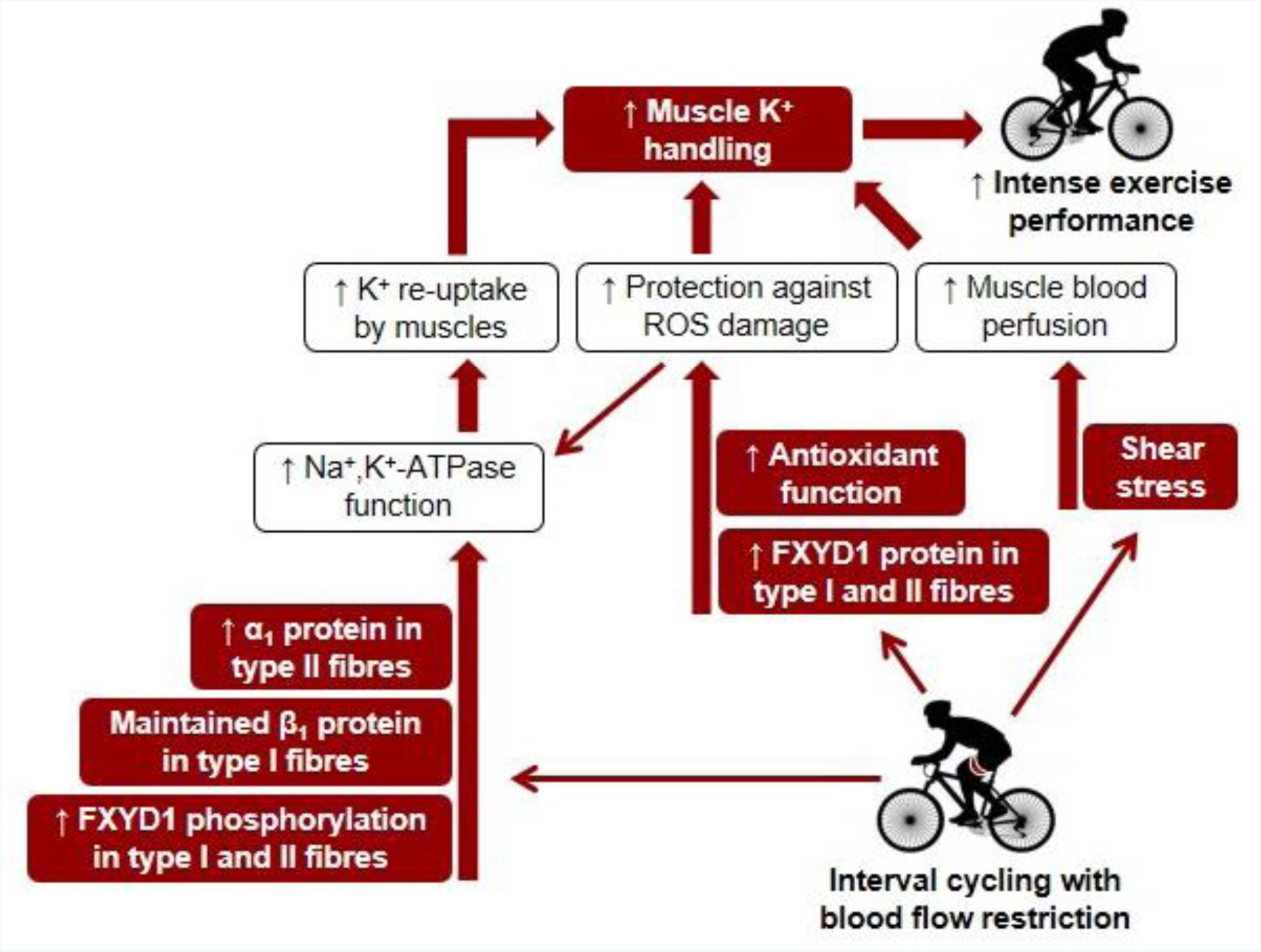
*Proposed factors underlying improvements in muscle K^+^ handling after training with blood flow restriction*. Based on the present outcomes, several factors likely contributed to improve skeletal muscle K^+^ handling after training with blood flow restriction (BFR), including fibre type-specific adaptations in abundance of Na^+^,K^+^-ATPase isoforms, increased protection against ROS, and elevated muscle blood flow.

## Supplemental tables

**Femoral arterial and venous plasma K+ concentration**

**Supplemental table 1.** *Mean femoral arterial and venous K^+^ concentration (mM) at rest before, during, and in recovery from single-leg, knee-extensor exercise at 25% of incremental peak power output before (Pre) and after (Post) six weeks of cycling without (CON) or with blood flow restriction (BFR) in men.*

|  | Rest | 20 s | 1 min | 3 min | 8 min | +20 s | +1 min | +2 min | +3 min |
| --- | --- | --- | --- | --- | --- | --- | --- | --- | --- |
| **CON PRE** |  |  |  |  |  |  |  |  |  |
| Arterial | 3.9±0.1 | 3.9±0.1 | 4.0±0.1 | 4.2±0.1 | 4.2±0.1 | 4.1±0.1 | 4.1±0.1 | 4.2±0.1 | 4.2±0.1 |
| Venous | 3.8±0.1 | 4.4±0.1 | 4.6±0.1 | 4.5±0.1 | 4.1±0.1 | 3.8±0.1 | 3.8±0.1 | 3.7±0.1 | 3.9±0.1 |
| **CON POST** |  |  |  |  |  |  |  |  |  |
| Arterial | 3.8±0.1 | 3.9±0.1 | 4.1±0.1 | 4.2±0.0 | 4.1±0.0 | 4.0±0.1 | 4.4±0.3 | 4.1±0.1 | 4.0±0.1 |
| Venous | 3.8±0.1 | 4.4±0.1 | 4.4±0.1(*) | 4.2±0.1* | 4.2±0.1 | 3.7±0.1 | 3.6±0.1 | 3.7±0.1 | 3.7±0.1 |
| **BFR PRE** |  |  |  |  |  |  |  |  |  |
| Arterial | 3.8±0.1 | 4.0±0.1 | 4.3±0.1 | 4.3±0.1 | 4.2±0.1 | 4.2±0.1 | 4.3±0.1 | 4.3±0.1 | 4.3±0.2 |
| Venous | 3.9±0.1 | 4.6±0.2 | 4.7±0.2 | 4.6±0.2 | 4.3±0.2 | 4.0±0.1 | 3.9±0.2 | 4.1±0.1 | 4.1±0.1 |
| **BFR POST** |  |  |  |  |  |  |  |  |  |
| Arterial | 3.9±0.1 | 4.0±0.1 | 4.2±0.0 | 4.2±0.1 | 4.2±0.1 | 4.1±0.1 | 4.2±0.0 | 4.1±0.0 | 4.1±0.1 |
| Venous | 3.8±0.1 | 4.4±0.1 | 4.5±0.1(*) | 4.3±0.1* | 4.2±0.1 | 3.8±0.1 | 3.7±0.1 | 3.8±0.1* | 3.9±0.1*# |

Data are means ± SE; n=8. \* $p < 0.05$ , different from PRE. (\*) $p < 0.1$ , tendency to be different from PRE. # $p <$
0.05, difference between CON and BFR in change from PRE to POST. (#) $p < 0.1$ , tendency for a difference between CON and BFR for the change from PRE to POST. Two two-way RM ANOVA were performed with sample time point and time (using raw values), or condition and time (using delta-sample data, i.e. POST-PRE) as factors.

**Supplemental table 2.**
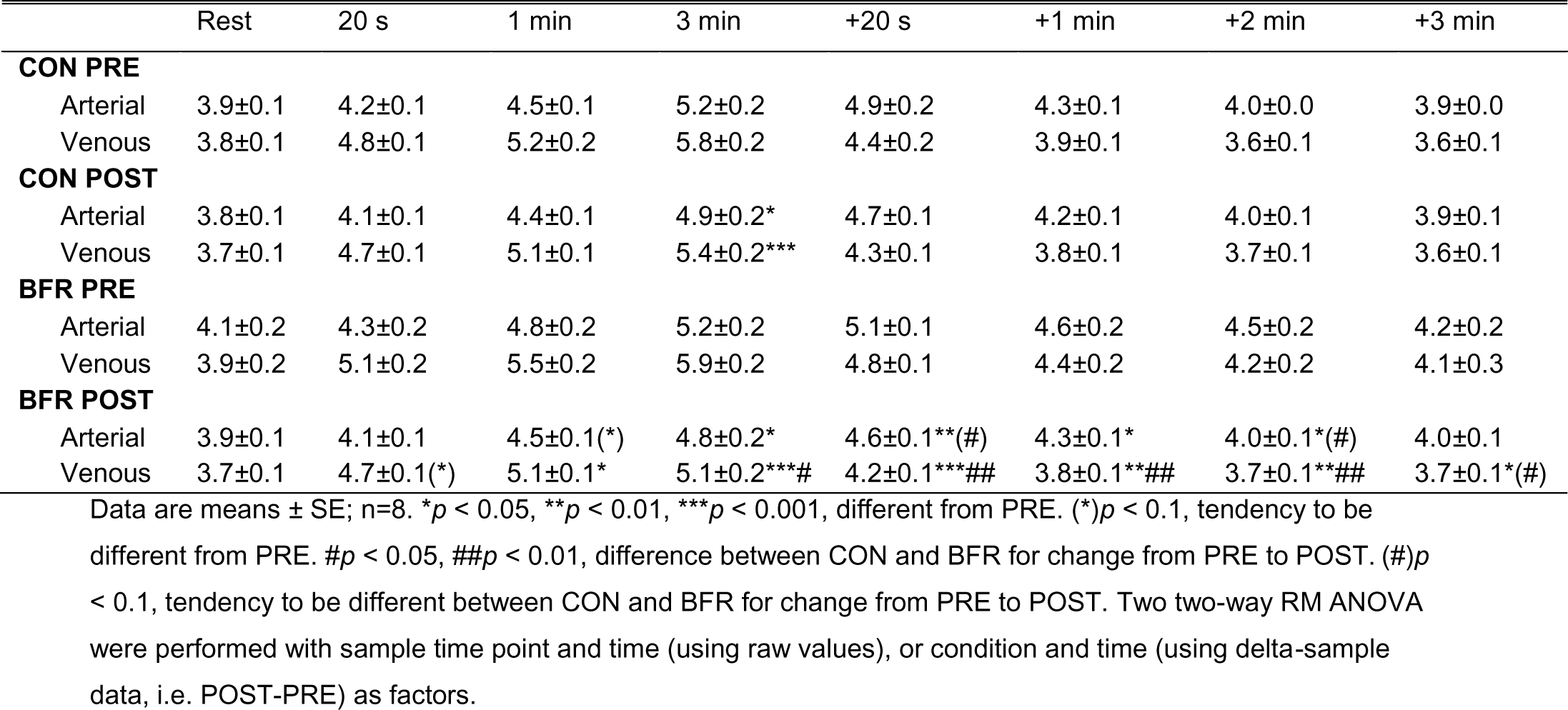
*Mean femoral arterial and venous K^+^ concentration (mM) at rest before, during, and in recovery from single-leg, knee-extensor exercise at 90% of incremental peak power output before (Pre) and after (Post) six weeks of cycling without (CON) or with blood flow restriction (BFR) in men.*

## Additional information

## Author contributions

Experimental procedures and analyses were performed at the Department of Nutrition, Exercise and Sport (NEXS), University of Copenhagen, Denmark. Training sessions were performed at a local gym (Fitnessdk Adelgade, Copenhagen, Denmark). DC and JB conceived the study. DC, MH, and JB designed experiments. DC, KE, VR, HV, MT, MN, TG, CS, ML, DB, and MH contributed to data collection and/or analysis. DC drafted the manuscript. DC, MH, and JB contributed to critically revising of this manuscript. All authors approved the final version.

## Acknowledgements

We thank Jens Jung Nielsen for excellent technical assistance with the experiments, and Anne K. Lund for the opportunity to train our participants at the training facility. We also thank the participants for their efforts, donation of muscle tissue and for a great time during training and tests. Finally, we would like to thank the students Peter Munck Sørensen, Tobias Buk Jørgensen, Daniel Lindsten, Eli Nolsøe Leifsson, Jesper Kirkebjerg, and Casper Bjerre Olsen for their efforts during training and experiments.

